# Multi-modal digital pathology for colorectal cancer diagnosis by high-plex immunofluorescence imaging and traditional histology of the same tissue section

**DOI:** 10.1101/2022.09.28.509927

**Authors:** Jia-Ren Lin, Yu-An Chen, Daniel Campton, Jeremy Cooper, Shannon Coy, Clarence Yapp, Juliann B. Tefft, Erin McCarty, Keith L. Ligon, Scott J. Rodig, Steven Reese, Tad George, Sandro Santagata, Peter K. Sorger

## Abstract

Precision medicine is critically dependent on better methods for diagnosing and staging disease and predicting drug response. Histopathology using Hematoxylin and Eosin (H&E) stained tissue - not genomics – remains the primary diagnostic method in cancer. Recently developed highly-multiplexed tissue imaging methods promise to enhance research studies and clinical practice with precise, spatially-resolved, single-cell data. Here we describe the “Orion” platform for collecting and analyzing H&E and high-plex immunofluorescence (IF) images from the same cells in a whole-slide format suitable for diagnosis. Using a retrospective cohort of 74 colorectal cancer resections, we show that IF and H&E images provide human experts and machine learning algorithms with complementary information that can be used to generate interpretable, multiplexed image-based models predictive of progression-free survival. Combining models of immune infiltration and tumor-intrinsic features achieves a hazard ratio of ∼0.05, demonstrating the ability of multi-modal Orion imaging to generate high-performance biomarkers.

## INTRODUCTION

The microanatomy of fixed and stained tissues has been studied using light microscopy for over two centuries^1, 2^, and immunohistochemistry (IHC) has been in widespread use for 50 years^3^. Histopathology review of hematoxylin and eosin (H&E) stained tissue sections, complemented by IHC and exome sequencing, remains the primary approach for diagnosing and managing many diseases, particularly cancer^4^. More recently, machine learning and artificial intelligence (ML/AI) approaches have been developed to automatically extract information from H&E images^5^, leading to rapid progress in computer-assisted diagnosis^6^. However, the H&E and IHC images used in existing digital pathology systems generally lack the precision and depth of molecular information needed to optimally predict outcomes, guide the selection of targeted therapies, and enable research into mechanisms of disease^7^.

The transition of histopathology from human inspection of physical slides to digital pathology^8^ is occurring concurrently with the introduction of methods for obtaining 10-100-plex imaging data from fixed tissue sections in a research setting (e.g., MxIF, CyCIF, CODEX, 4i, mIHC, MIBI, IBEX, and IMC)^9–15^. These high-plex imaging approaches enable deep morphological and molecular analysis of normal and diseased tissues from humans and animal models^12, 16–19^ and generate spatially resolved information that is an ideal complement to other single cell methods, such as scRNA sequencing. Whereas some imaging methods require frozen samples, those that are compatible with formaldehyde-fixed and paraffin-embedded (FFPE) specimens – the type of specimens universally acquired for diagnostic purposes – make it possible to tap into large archives of human biopsy and resection specimens^20, 21^. Many high-plex imaging studies performed to date on human cohorts involve tissue microarrays (TMAs; arrays of many 0.3 to 1 mm specimens on a single slide) or the small fields of view characteristic of mass-spectrometry based imaging^9, 11^. However, whole-slide imaging is required for clinical research and for diagnosis, both to achieve sufficient statistical power^22^ and as an FDA requirement^23^.

Histopathology review of H&E images is a top-down approach in which human experts draw on prior knowledge about the abundances and morphologies of cellular and acellular structures prognostic of disease or predictive of drug response. This prior knowledge, summarized in resources such as the American Joint Committee on Cancer’s staging manual^24^, is based on thousands of clinical research papers and numerous clinical trials. In contrast, research using highly multiplexed imaging most commonly involves a bottom-up approach in which cell types are enumerated and an attempt is made to identify single-cell features or cell neighborhoods associated with disease, for example using spatial statistics^9, 11^. These types of high-plex imaging studies are in their infancy and have not yet been subjected to rigorous validation in a clinical setting. Thus, a substantial opportunity exists to link established histological workflows with emerging multiplexed methods in research and diagnostic settings, thereby allowing deep knowledge of tissue anatomy (from H&E images)^25^ to be combined with precise single cell-data on tumors and their microenvironment.

We reasoned that an ideal instrument for bridging top-down and bottom-up approaches would perform whole-slide imaging (WSI),^26^ have sufficient plex and resolution to distinguish tumor, immune and stromal cell types, and enable reliable and efficient data acquisition with minimal human intervention. In current practice, combining high-plex immunofluorescence and H&E imaging requires the use of different tissue sections^27^. However, collection of same-cell multi-modal images would have the substantial advantage of enabling one-to-one comparison of cell morphologies and molecular properties. Same-cell H&E and high-plex imaging would also facilitate computational approaches that combine single-cell molecular profiling with rapid developments in the use of ML/AI to interpret H&E data^28^.

The relative complexity of existing highly multiplexed imaging assays has prevented their widespread adoption in the clinic; the current standard in clinical research is 5 to 6-plex imaging of tissue sections using a Perkin Elmer Vectra Polaris™ (now Akoya PhenoImager HT™)^29, 30^. However, a first-principles analysis suggests that a minimum of 16-20 molecular (IF) channels are required for tumor profiling (**Extended Data Table 1**): 10-12 to subtype major immune cell types (blue), 2-3 to detect and subtype tumor cells and states (green), 2-4 to identify relevant tissue structures (yellow),1-3 to examine tumor cells states or therapeutic mechanisms (grey), plus a nuclear stain to locate cell nuclei (pink). We reasoned that achieving this or higher plex in a diagnostic or high-throughput research setting would require acquisition of many fluorescent channels in parallel (one-shot imaging) rather than the sequential process developed by Gerdes et al.^10^ and subsequently extended by our group^15^ and others^31^.

In this paper, we describe the development of an approach for one-shot, whole-slide, 16 to 18- channel immunofluorescence (IF) imaging, followed by H&E staining and imaging of the same cells. Using FFPE specimens from multiple tumor types, we also compare the performance of the “Orion™” approach and a commercial-grade instrument that implements it, with established IHC and cyclic data acquisition by CyCIF^32^. We find that joint analysis of H&E and IF same-section Orion images substantially improves our ability to identify and interpret image features by facilitating the transfer of anatomical annotation from H&E images to high-plex data (e.g., by distinguishing normal tissue from a tumor) and also the other way round (e.g., by enabling subtyping of immune cells that are indistinguishable in H&E data). We show that machine learning (ML) models generated from molecular analysis of high-plex IF images can be combined with ML of H&E images to aid in feature identification and interpretation (substantially extending previous data on joint analysis of molecular and H&E images)^33, 34^. In a proof of principle study, we use whole-slide Orion imaging to identify spatial biomarkers prognostic of tumor progression in independent 30-40-patient human colorectal cancer (CRC) cohorts (n = 74 patients total). A combination of top-down and bottom-up methods enabled the generation of biomarker-based models with Hazard Ratios of 0.05 to 0.15. The Orion method is scalable to the large multi-center cohorts needed to test and validate these proof of principle biomarkers for eventual use in patient care.

## RESULTS

### Constructing and testing the Orion platform

We investigated multiple approaches for achieving one-shot high-plex IF followed by H&E imaging of the same cells (i.e., from the same tissue section). Overlap in the excitation and emission spectra of most widely used fluorophores limits the number of separable fluorescence channels (typically to five to six) that can be accommodated within the wavelengths useful for antibody labeling (∼350 to 800 nm). This can be overcome using tuned emission and excitation filters and spectral deconvolution (e.g., of 6 - 10 channels)^35^ or by dispersing emitted light using a diffraction grating and then performing linear unmixing^36, 37^. However, unmixing of complex spectra (e.g., from a tissue image stained with 10 or more fluorophores) has historically resulted in a substantial reduction in sensitivity and has not been widely implemented. Simultaneous high-plex imaging of tissue specimens therefore required innovation in the optical platform as well as careful selection of fluorophores.

With support from an NCI SBIR grant, a commercial-grade Orion instrument was developed. The instrument utilizes seven lasers (**Fig. 1a** and **Extended Data Fig. 1**) to illuminate the sample and collect emitted light with 4X to 40X objective lenses (0.2 NA to 0.95 NA; Orion data is this paper were collected with a 20X 0.75 NA objective) followed by multiple tunable optical filters^38^ that use a non-orthogonal angle of incidence on thin-film interference filters to shift the emission bandpass^39^. These filters have 90-95% transmission efficiency and enable collection of 10 - 15 nm bandpass channels with 1 nm center wavelength (CWL) tuning resolution over a wide range of wavelengths (425 to 895 nm). Narrow bandpass emission channels improve specificity but substantially reduce signal strength; we overcame this problem by using excitation lasers that are ∼10 times brighter than conventional LED illuminators and by using a sensitive scientific CMOS detector (camera). Raw image files were then processed computationally to correct for system aberrations such as geometric distortions and camera non-linearity^40^, followed by spectral extraction to remove crosstalk and isolate individual fluorophore signals (and thus, the antibodies to which they were conjugated). The features of single cells and regions of tissue were then computed using MCMICRO software^41^.

**Fig. 1.**
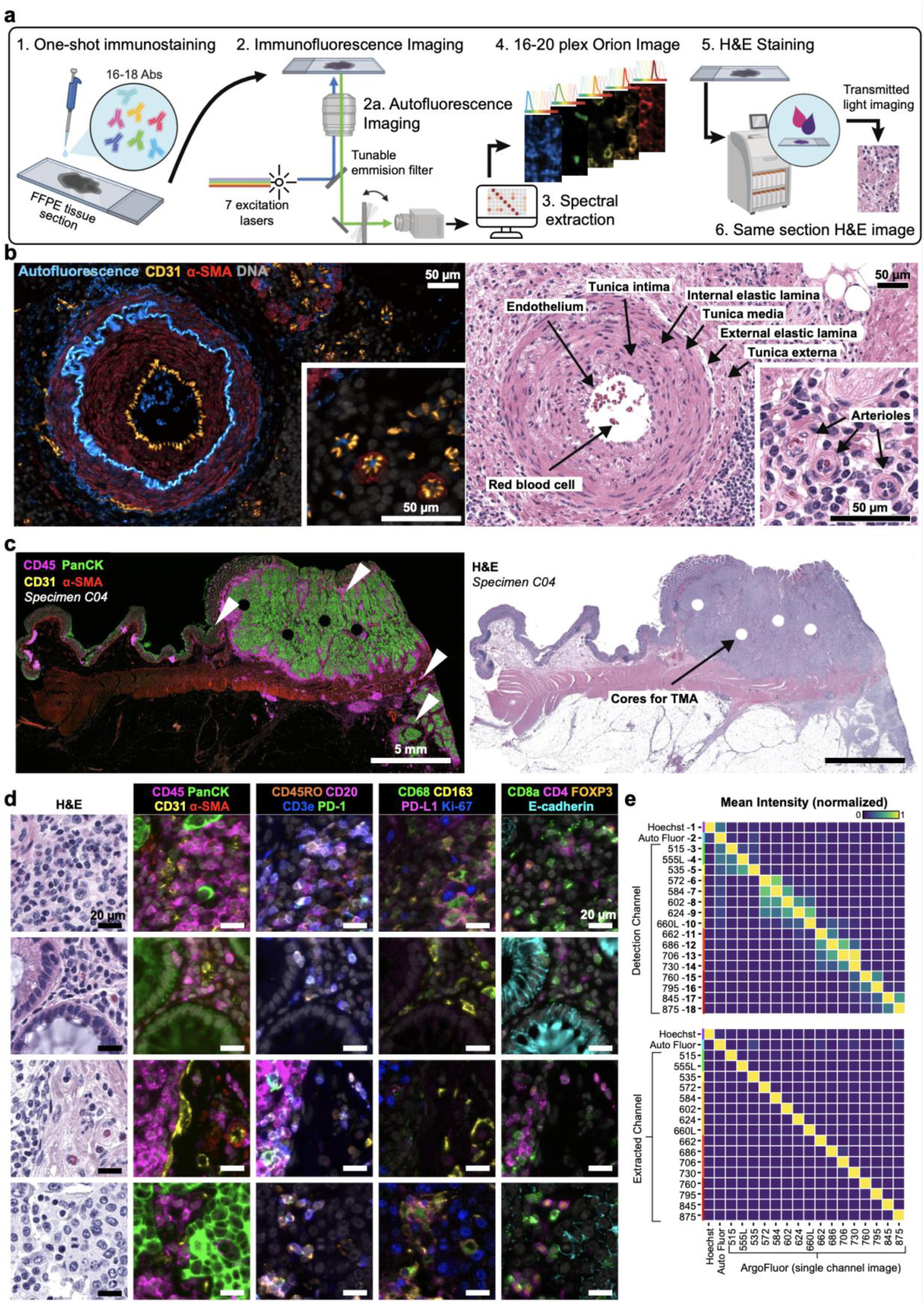
Same-section immunofluorescence and H&E using the Orion™ Platform. **a,** Schematic of one-shot 16 to 20-channel multiplexed immunofluorescence imaging with the Orion™ method followed by Hematoxylin and Eosin (H&E) staining of the same section using an automated slide stainer and scanning of the H&E-stained slide in transillumination (brightfield) mode. This method of discriminating the emission spectra of fluorophores is repeated using seven excitation lasers spaced across the spectrum (see **Extended Data Fig. 1a** and Methods section). Using polychroic mirrors and tunable optical filters, emission spectra are extracted to discriminate up 20 channels including signal from fluorophore-labelled antibodies (15-19 in most experiments), the nuclear stain Hoechst 33342, and tissue intrinsic autofluorescence. **b,** Left panels: Orion multiplexed immunofluorescence image showing CD31, α-SMA, Hoechst (DNA), and signal from the tissue autofluorescence channel (AF) from a colorectal cancer FFPE specimen (C04); this highlights an artery outside of the tumor region with red blood cells in the vessel lumen and elastic fibers in the internal and external elastic lamina of the vessel wall, numerous smaller vessels (arterioles), and stromal collagen fibers (inset displays arterioles). Right panels: images of the H&E staining from the same tissue section (histologic landmarks are indicated). Scalebars 50 µm. **c,** Orion multiplexed immunofluorescence image (showing CD45, pan-cytokeratin, CD31, and α-SMA) from a whole tissue FFPE section of a colorectal cancer (C04) and matched H&E from the same section. Holes in the images are regions of tissue (‘cores’) removed in the construction of TMAs. Scalebar 5 mm. **d,** Zoom-in views of the regions indicated by arrowheads in panel **c**; marker combinations indicated. Scalebars 20 µm. **e,** Intensities of fluorochromes (columns in heatmaps) in each Orion channel (rows in heatmaps) prior to (top) and after (bottom) spectral extraction. The extraction matrix was determined from control samples scanned using the same acquisition settings that were used for the full panel. The control samples included: unstained lung tissue (for the autofluorescence channel), tonsil tissue stained with Hoechst, and tonsil tissue stained in single-plex with ArgoFluor-conjugates used in the panel (for the biomarker channels). The values in each column were normalized to the maximum value in the column.

We tested >100 chemical fluorophores from different sources and identified 18 ArgoFluor™ dyes that were compatible with spectral extraction enabled by discrete sampling. Key criteria were: (i) emission in the 500 - 875 nm range; (ii) high quantum-efficiency; (iii) good photostability; and (iv) compatibility with each other in high-plex panels (**Extended Data Fig. 1a, Extended Data Table 1& 2**). ArgoFluor dyes were covalently coupled to commercial antibodies directed against lineage markers of immune (e.g., CD4, CD8, CD68), epithelial (cytokeratin, E-cadherin), and endothelial (CD31) cells as well as immune checkpoint regulators (PD-1, PD-L1), and cell state markers (Ki-67), to generate panels suitable for studying the microenvironment and architecture of epithelial tumors and adjacent normal tissue (**Extended Data Fig. 1b;** the logic underlying Orion panels is show in **Extended Data Fig. 2a**). An accelerated aging test demonstrated excellent reagent stability, estimated to be >5yr at - 20°C storage (**Extended Data Fig. 1c)**.

Because eosin fluoresces strongly in the 530 - 620 nm range, it proved impractical to perform H&E staining prior to IF (although alternatives to H&E compatible with IF have been described)^42^. However, H&E images could be obtained after one or a small number of IF cycles when staining was performed using an industry-standard Ventana automated slide stainer (or similar machines from other vendors)^43^. No established standard exists for evaluating the quality of these or other digital H&E images^44^ and comparison across methods is complicated by variation in H&E color intensity among even clinical histopathology centers^45^. We therefore showed four practicing pathologists images of tissue sections that had been subjected to one or more IF staining cycles followed by fluorophore bleaching and asked whether practitioners could distinguish these images from serial section controls that had been stained with H&E in the standard manner in a clinical facility. The Orion instrument has an integrated brightfield mode, but the H&E images used in this study were also acquired using an Aperio GT450 microscope (Leica Biosystems), which is a gold standard for diagnostic applications and facilitated image comparison by human experts.^46^ (**Fig. 1a**). When our panel of pathologists compared control H&E images with those obtained after by Orion, they found them to be indistinguishable and “diagnostic grade” (**Extended Data Fig. 1f**).

### Validating high-plex one-Shot fluorescence imaging

To test the Orion approach, three types of data were collected: (i) whole slide images of both human tonsil, a standard tissue for antibody qualification, and human lung cancer, a particularly common cancer type; (ii) images of a TMA that contains 30 different types of normal, non-neoplastic disease, and tumor samples from 18 tissue types, including brain, breast, colon, kidney, liver, lung, lymph node, ovary, pancreas, prostate, skin, small intestine, spleen, testis, and tonsil (iii); whole-slide images of 74 stage I-IV colorectal cancer (CRC) resections obtained from the archives of the Brigham and Women’s Hospital Pathology Department (these resections were split into two cohorts with 40 and 34 patients each as indicated in **Extended Data Table 3**). We tested and optimized the antibody panel on tonsil tissue and then applied it successfully to the lung cancer specimen (**Extended Data Fig. 2b**), TMA (**Extended Data Fig. 2c**), and CRC cohort. We also collected data from a dedicated autofluorescence channel (445 nm excitation / 485 nm emission, CWL) both to extract natural fluorescence from the IF channels and improve biomarker signal to noise ratio (SNR), and to provide information on naturally fluorescent structures such as connective tissues and components of blood vessels (**Fig. 1b**)^47^. In each case, we performed 18 - 20 plex imaging (16-18 antibody channels, autofluorescence and a nuclear stain) plus H&E. However, exploratory studies suggest that it should be possible to add 2-4 additional antibody channels to the method following further optimization of fluorophores and optical systems (see Methods).

In whole slide images of lung, tonsil, and CRC, inspection of extracted images revealed error-free whole-slide imaging of 1,000 or more adjacent tiles (area up to 35 by 20 mm; **Fig. 1c**)^48^ as well as bright in-focus staining of cellular and cellular substructures within each tile (**Fig. 1d**). To quantify the effectiveness of spectral extraction, we imaged serial sections of human tonsil tissue each stained with a single antibody conjugated to a different ArgoFluor and then recorded data in all channels. Under these conditions, cross talk between adjacent channels averaged ∼35%. Spectral extraction reduced this to <1% (in all but a few cases crosstalk among all pairs of channels was <0.5%; **Fig. 1e**). As a result, when a tissue section was subjected to multiplexed antibody labeling, we observed correlated signals only for antibodies that stain targets co-localized on the same types of cells (e.g., co-staining of T-cell membranes by antibodies against CD3e and CD4 resulted in correlation of the corresponding fluorescence channels; **Extended Data Fig. 1e**).

The staining patterns observed with ArgoFluor-antibody conjugates were similar to those obtained by conventional IHC performed on the same specimen using the same antibody clones (as described in Du et al.^49^, one-to-one comparison of IF and IHC is not possible given fundamental differences in imaging modalities; **Fig. 2a** and **Extended Data Fig. 3a**). We also compared Orion data to data acquired from a serial tissue section using the well-established cyclic immunofluorescence (CyCIF) method^15^. We found that the images looked very similar and that fractions of cells scoring positive for the same markers across the two methods were highly correlated (**Fig. 2b, 2c** shows four examples with ρ = 0.8 to 0.9). However, when marker positive cells were less abundant, cell counts were subject to greater statistical fluctuation from one serial sections to the next, and data from CyCIF and Orion were less correlated (e.g. ρ = 0.55 for FOXP3 positivity; **Extended Data Fig. 3b**) . Nonetheless, projections of high dimensional Orion data using t-SNE successfully resolved multiple immune and tumor cell types (**Fig. 2d** and **Extended Data Fig. 3c**).

**Fig. 2.**
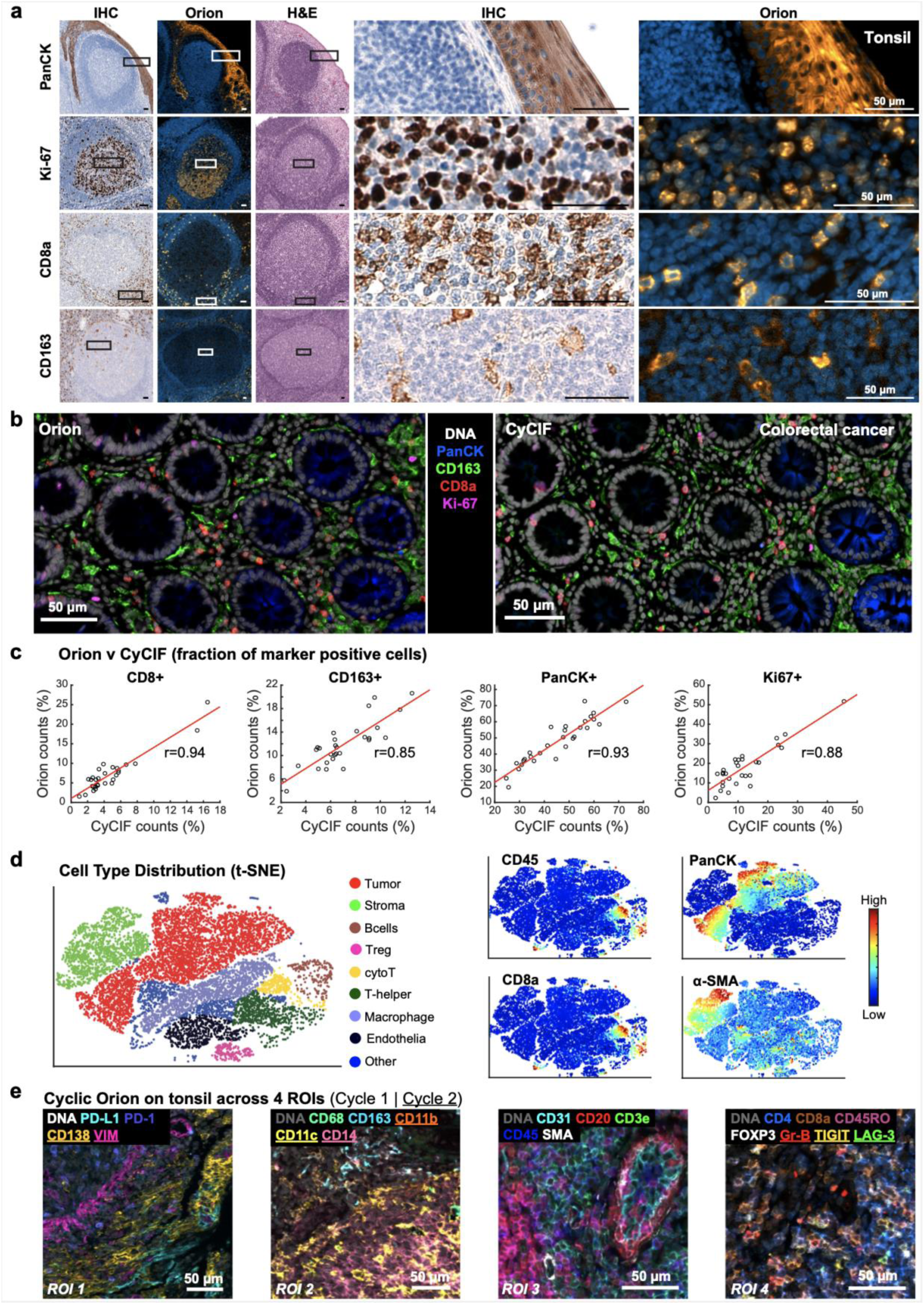
Qualifying 16-plex single-shot Orion antibody panel. **a,** Panels of images from FFPE tonsil sections showing single-antibody immunohistochemistry (IHC) for pan-cytokeratin, Ki-67, CD8a, CD163, and the matching channels extracted from 16-plex Orion immunofluorescence (IF) images (H&E stain was performed on the same section as the Orion imaging). Scalebars 50 µm. **b,** Orion IF images and cyclic immunofluorescence (CyCIF) images from neighboring sections of an FFPE colorectal adenocarcinoma; Scalebars 50 µm. The CyCIF images collected using 2×2 binning while Orion images were obtained with no binning. **c,** Plots of the fraction of cells positive for the indicated markers from whole slide Orion IF and CyCIF images acquired from neighboring sections from 29 FFPE colorectal cancer specimens. Pearson correlation coefficients are indicated. **d,** t-distributed stochastic neighbor embedding (t-SNE) plots of cells derived from CyCIF (left panels) and Orion IF images (right panels) of a FFPE colorectal cancer specimen (C01) with the fluorescence intensities of immune (CD45, pan-cytokeratin, CD8a, α-SMA) markers overlaid on the plots as heat maps. **e,** Orion images of FFPE tonsil tissue showing antibodies imaged across two cycles. 23 of 29 antibodies are displayed across four marker groups from four different regions of interest (labeled ROI 1-4). Markers from cycle 2 are underlined. The locations of the four ROIs in the whole slide image are shown in **Extended Data Fig. 5a**). Scalebars 50 µm.

To test the repeatability of the method, sample processing and imaging of CRC Cohort 1 (n = 40 specimens) was performed at RareCyte, and processing and imaging of Cohort 2 (n = 34 different specimens) was performed at HMS on a different instrument by different operators; six specimens from Cohort 1 were imaged at both RareCyte and HMS. Corresponding pairs of images from these six specimens looked very similar and when cell count data from all 12 images was subjected to unsupervised clustering, batch effects were not observed (**Extended Data Fig. 4a-c**). Thus, the Orion method generates results that are qualitatively similar to those obtained using conventional IHC and quantitative marker intensities are similar between Orion and CyCIF.

There are situations in which data from 16-20 fluorescent channels is likely to be insufficient for identifying cell types of interest. We therefore asked whether multiple rounds of Orion data collection could be performed on the same cells using a cyclic approach^10, 15^. We stained tonsil tissue with 16 ArgoFluor-conjugated antibodies and collected IF data plus autofluorescence and an image of DNA in the Hoechst channel. Slides were subjected to oxidation with hydrogen peroxide (bleaching), stained with 13 additional antibodies (this number was based on reagent availability), followed by data acquisition and processing for H&E and brightfield imaging. We found that crisp, high SNR second-round images could be obtained using a cyclic approach, yielding a 32-plex Orion image (if same-cell H&E is included; **Fig. 2e** and **Extended Data Fig. 5a**). We confirmed that the inter-cycle bleaching step reduced ArgoFluor intensity by >95% and that crosstalk from one cycle to the next was therefore low (**Extended Data Fig. 5b**). We also found that it was possible to perform multiple rounds of CyCIF after one round of Orion (**Extended Data Fig. 5c**). Moreover, although many cycles of IF staining and bleaching reduced H&E image quality, our pathology team judged H&E images collected after two IF and photobleaching steps to indistinguishable from controls and therefore diagnostic grade (**Extended Data Fig. 5d, e**). We conclude that two-cycle Orion imaging retains IF and H&E image quality, opening the door to efficient 32-36 plex multi-modal imaging. Exploratory studies suggest room for further development of cyclic and high-plex Orion imaging although more rigorous approaches to scoring H&E image quality will be required.

### Integrated analysis of IF and H&E images

When same-cell H&E and IF data were compared, we found that molecular labels obtained from IF enabled more complete enumeration of lymphocytes than inspection of H&E images by trained pathologists alone; for example, CD4 and CD8 T cell and B cell lineages look similar by H&E but clearly distinguishable by IF (arrows in **Fig. 3a**). We also identified many cell types and cell states that were more readily defined in H&E images based on morphologic features than by immunofluorescence staining; this included cells such as eosinophils and neutrophils whose morphology is highly characteristic but which had no lineage markers in our Orion panels, as well as the prophase, metaphase, anaphase and telophase stages of mitosis (arrows and dashed lines in **Fig. 3b**). A wide variety of acellular structures such as basement membranes, mucin pools, necrotic domains, etc. were also more readily scored in H&E than IF images. To begin to quantify the amount of complementary information in H&E and IF images, we computed the fraction of all cells (as identified by nuclear segmentation) in the 40-specimen CRC Cohort 1 that could not be assigned a clear identity using IF images; we found that this varied from 6.5 to 42% of total nuclei (median = 16%) (**Fig. 3c**). We have previously observed a similar fraction of “unidentifiable” cells following 40-60 plex CyCIF imaging^22^ and surmised that these cells were either negative for all antibody markers included in the panel or had morphologies that are difficult to segment^50^.

**Fig. 3.**
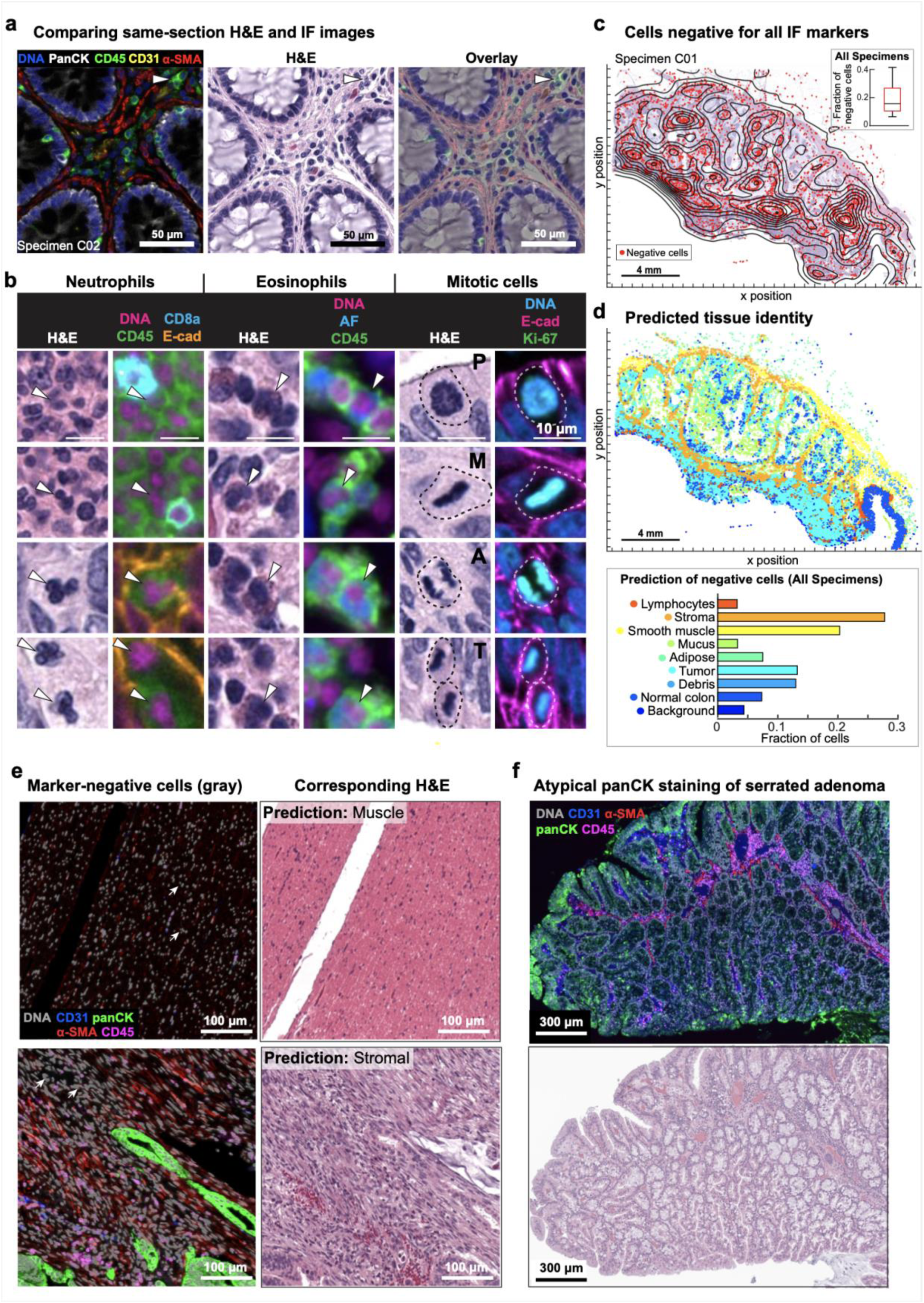
Combined H&E and Orion to identify cell/tissue types. **a,** Representative images of Orion IF and same-section H&E from an area of normal colon (from colorectal cancer resection specimen C02). Scalebars 50 µm. **b,** Cell types not specifically identified by markers in the Orion panel but readily recognized in H&E images including neutrophils, eosinophils, and cells undergoing mitoses (selected cells of each type denoted by arrowheads and dashed lines). Scalebars 10 µm. **c,** Spatial maps of the positions of cells (∼15% of total cells) that were not detected by the Orion IF panel in a colorectal cancer specimen overlaid onto the corresponding H&E image (specimen: C01); red dots denote cells with identifiable nucleus but not subtyped using the antibody panel. **d,** Upper panel: Spatial map of nine tissue classes determined from the H&E image using a convolutional neural network (CNN) model for various cell types as indicated^51^. Lower panel: Percent of total of “unidentifiable” (negative) cells assigned to a specific tissue class by the CNN applied to the H&E image. **e,** Example same-section Orion IF and H&E images from areas enriched for ‘non-detected’ cells; examples include areas predicted to be rich in stroma and smooth muscle; Scalebars 100 µm. **f,** Orion IF and H&E images from colorectal cancer resection specimen C26, showing an area of serrated adenoma with low pan-cytokeratin expression (markers as indicated). Whole slide image indicating the location of this region is shown in **Extended Data Fig. 5f**. Scalebars 300 µm.

To identify cells missing labels in Orion IF data, we used a previously published ML model trained on H&E images^51^ (see Methods for details of this model and its performance). We found that >50% were predicted to be smooth muscle, stromal fibroblasts or adipocytes (**Fig. 3d**); these predictions were confirmed by visual inspection of the H&E images by pathologists (**Fig. 3e**). We also examined specimens (e.g., from patient 26, **Fig. 3f** and **Extended Data Fig. 5f**) in which a subset of epithelium was difficult to identify by IF because it was weakly stained by pan-cytokeratin, E-cadherin, and immune markers. Inspection of H&E images showed that these weakly-staining cells corresponded to a serrated adenoma that was distinct from nearby domain of invasive low-grade adenocarcinoma in which tumor cells stained strongly for pan-cytokeratin and E-cadherin. Differential staining of cytokeratin isoforms in serrated adenoma and adenocarcinoma has been described previously^52^ and we speculate that in cases such as specimen C26, it reflects clonal heterogeneity. Regardless, low staining intensity interferes with IF-based cell type calling for a large fraction of the tumor cells in the specimen. From these findings we conclude that the availability of H&E and IF images of the same set of cells substantially increases the fraction of cell types and states that can be identified as compared to either type of data alone. This is particularly true of cell types for which specific molecular markers do not exist (e.g., stromal fibroblasts) or are not included in the panel (e.g., neutrophils) and markers that are lost due to tumor sub-clonality (e.g., specific cytokeratin isoforms). Cells that are highly elongated or have multiple nuclei and are difficult to segment (e.g., muscle cells) are also commonly lost to computational analysis of IF data but highly distinctive in H&E images.

### Identifying tumor features predictive of disease progression

The classification of cancers for diagnostic purposes using American Joint Committee on Cancer (AJCC/UICC-TNM classification) criteria is based primarily on tumor-intrinsic characteristics (tumor, lymph node, and metastases, the TNM staging system)^53^. However, the extent and type of immune infiltration also plays a major role in therapeutic response and survival^54^. In colorectal cancer (CRC) this has given rise to a clinical test, the Immunoscore^®55^, that quantifies features of the intratumoral and tumor-proximal immune response to predict CRC progression as measured by progression-free survival (PFS) or overall survival (OS). The Immunoscore has been validated in multicenter cohort studies and shown to predict time to recurrence in stage III cancers in a Phase 3 clinical trial^56^. The Immunoscore uses IHC to evaluate the number of CD3 and CD8-positive T cells at the tumor center (CT) and the invasive margin (IM; for Immunoscore this is defined as a region encompassing 360 μm on either side of the invasive boundary; in our work we set this to ± 100 μm from the boundary). The hazard ratio (HR; the difference in the rate of progression) between patients with tumors containing few immune cells in both the CT and the IM (Immunoscore = 0) and patients with tumors containing many cells in both compartments (Immunoscore = 4) has been reported to be 0.20, (95% CI 0.10–0.38; p < 10^-4^) in a Cox regression model, with increasing score correlating with longer survival^59^. This is a clinically significant difference that can be used to inform key treatment decisions: for example, whether or not to deliver adjuvant therapy (chemotherapy after surgery)^60^. Because chemotherapy is associated with significant adverse effects, requires infusion or injection in a healthcare setting, and is expensive, it is highly desirable that patients who are unlikely to experience disease recurrence be spared the burden of adjuvant therapy.

Using Orion data, we developed software scripts to recapitulate key aspects of the Immunoscore using Progression Free Survival (PFS) as an outcome measure. First, we detected the tumor-stromal interface and generated masks that matched the criteria for CT and IM (± 100 μm around the tumor boundary; **Fig. 4a**). CD3 and CD8 positivity in single cells was determined by Gaussian Mixture Modeling^61^ with the median positive fraction for each marker (CD3 or CD8) in each region (CT or IM) across all 40 CRC cases used as the cutoff for assigning a subscore of 0 or 1; the sum of the four subscores was used as the final score for Image Feature Model 1 (IFM1; **Fig. 4b**). Parameters for computing IFM1 such as the size of the invasive margin and the staining threshold for scoring cells positive and negative were set *a priori* (naively) without any parameter tuning to reduce the risk of over-training; IFM1 nonetheless yielded a hazard ratio similar to Immunoscore itself on Cohort 1 (HR = 0.14; 95% CI 0.06-0.30; p = 7.63 × 10^-5^) (**Fig. 4c**), Next, we used the underlying logic of Immunoscore to leverage multiple Orion channels. A total of 13 immune focused markers were used to generate ∼15,000 marker combinations (IFMs), each composed of four markers within the CT and IM domains (**Fig. 4d**). Scores for each CRC case were binarized into high and low scores based on median intensities (again, without any parameter tuning). When HRs were calculated we found that nearly 600 IFMs exceeded IFM1 in performance (**Extended Data Fig. 6a-c**). The top 10 IFMs were insignificantly different from each other, and we chose one (IFM2) for further analysis; it exhibited an HR = 0.05 (95% CI: 0.02-0.10, p = 5.5 × 10^-6^) (**Fig. 4d** and **4e**) and comprised the fractions of α-SMA^+^ cells in the CT, and CD45^+^, PD-L1^+^, and CD4^+^ cells in the IM. Leave-one-out resampling showed that IFM2 was significantly better than IFM1 with respect to HR (adjusted p value based on the Benjamini-Hochberg Procedure p_adj_ = 7.3 × 10^-21^; **Fig. 4f, Extended Data Fig. 6d**). To determine whether this result could be generalized to other specimens, we tested the performance of an IFM2 model created using Cohort 1 on specimens in Cohort 2. Once again, we observed a statistically significant discrimination between progressing and non-progressing tumors (HR = 0.17; 95% CI: 0.05 to 0.56, p = 6.9 × 10^-3^; **Fig. 4g**). We conclude that multiplexed immunoprofiling data extracted from Orion images of CRC resections can be used to generate high performance prognostic biomarkers.

**Fig. 4.**
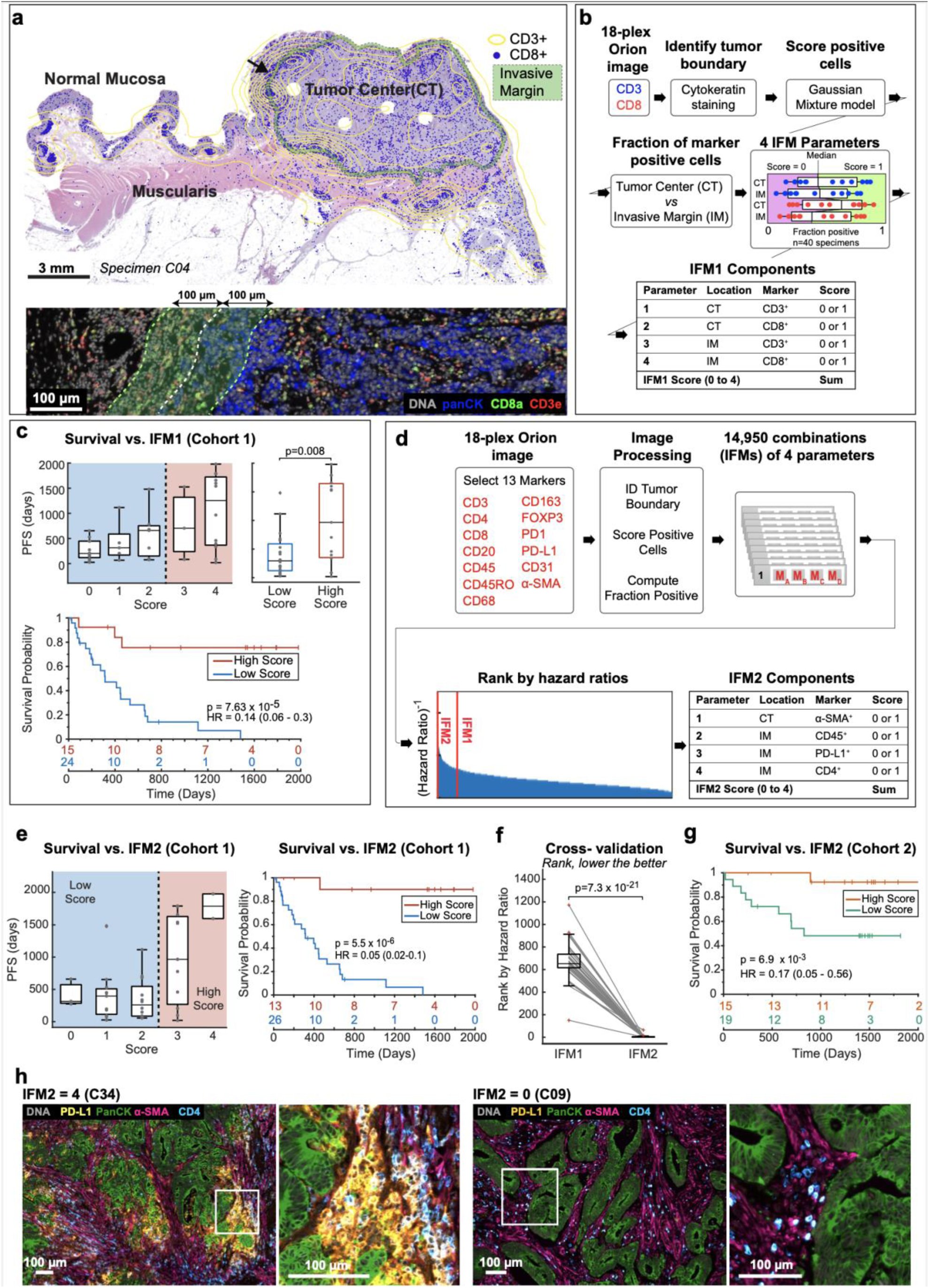
Recapitulating and extending the Immunoscore tissue immune test using Orion images. **a,** Map of tumor center and invasive-margin compartments for specimen C04 overlaid on an H&E image with the density of CD3^+^ cells shown as a contour map (yellow) and the positions of CD8^+^ T cells as blue dots. The arrow indicates the zoom-in images shown below. Lower panel shows selected channels from a portion of the Orion image for C04 spanning the invasive boundary (denoted by green shading). **b,** Flow chart for the calculation of Image Feature Model 1 (IFM1) that recapitulates key features of the Immunoscore test. **c,** Upper panel: Box-and-whisker plots for progression-free survival (PFS) for 40 CRC patients based on actual IFM1 scores (midline = median, box limits = Q1 (25th percentile)/Q3 (75th percentile), whiskers = 1.5 inter-quartile range (IQR), dots = outliers (>1.5IQR) or scores stratified into two classes as follows, low: score ≤ 2, high: score = 3 or 4 (pairwise two-tailed t-test p = 0.002. Lower panel: Kaplan Meier plots computed using IFM1 binary classes (HR, hazards ratio; 95% confidence interval; logrank p-value). **d,** Flow chart for calculation of additional models that use the underlying logic of Immunoscore but considering 13 markers. The image processing steps are the same as in panel *a*. The rank positions of IFM1 and IFM2 are shown relative to all other 14,950 combinations of parameters that were considered. **e,** (Left) Box-and-whisker plots for PFS for 40 CRC patients based on IFM2 scores, with ranges as defined in c. (Right) Kaplan Meier plots for Cohort 1 computed using IFM2 binary classes stratified into two classes as follows, low: score ≤ 2, high: score = 3 or 4 (HR, hazards ratio; 95% confidence interval; logrank p-value). **f.** Plot of leave-one-out cross-validation of ranks from IFM1 and IFM2 (unadjusted p = 4.9 × 10^-26^ and adjusted using the Benjamini-Hochberg Procedure; p=7.3 × 10^-21^); bootstrapping of hazard ratios is shown in **Extended Data Fig 6d.** Detailed analysis was described in the methods section and pairwise two-tailed t-test were used unless otherwise mentioned. **g,** Kaplan Meier plot for Cohort 2 computed using IFM2 binary classes stratified into two classes as follows, low: score ≤ 2, high: score = 3 or 4 (HR, hazards ratio; 95% confidence interval; logrank p-value). **h,** Representative Orion IF images of cases with high IFM2 (score = 4 in specimen C34) and low IFM2 (score = 0 in specimen C09). IF images show DNA, pan-cytokeratin, α-SMA, CD45, and PD-L1; Scalebars 100 µm.

Inspection of images from IFM2 tumors exhibiting slow progression (e.g., patient C34) revealed high-levels of PD-L1^+^ cells (**Fig. 4h**, yellow) adjacent to pan-cytokeratin positive tumor cells (green); based on overlap of PD-L1^+^,CD68 and CD45 staining we conclude that PD-L1^+^ cells are likely myeloid in origin, as described previously^22^. In C34, α-SMA stained tumor proximate stromal cells – most likely fibroblasts – were also well-infiltrated with CD4^+^ T cells. By contrast, in a patient with rapid progression (e.g., patient C09), PD-L1 levels were below the level of detection and CD4^+^ cells were less abundant in the stroma. By H&E, IFM2-high tumors exhibited extensive lymphohistiocytic chronic inflammation including large lymphoid aggregates and tertiary lymphoid structures (TLS) at the tumor invasive margin^62^, whereas IFM2-low tumors had relatively few lymphoid aggregates and no TLS (**Fig. 4h** and **Extended Data Fig. 6e**). Although IFM2-low tumors were also more invasive than IFM2-high tumors, IFM score was independent of histologic subtype (e.g., conventional vs. mucinous morphology) and did not correlate with histologic grade (low vs. high grade carcinoma). Thus, IFM2 is likely to capture activity of the immune microenvironment around the invasive tumor margin as well changes in tumor-associated fibroblasts. However, deeper phenotyping of more specimens will be required to identify which molecular features of IFM2 are important for predicting progression. However, we conclude that Orion data can be used to automate previously described image-based biomarkers based on single-channel IHC and identify new marker combinations that significantly outperform them (see limitations sections for further discussion of this point).

### Identifying new progression markers

As an unbiased bottom-up means of identifying new progression models, we used spatial Latent Dirichlet Allocation (Spatial-LDA)^63^. This approach is a modification of the LDA method developed for analysis of text^64^ that enables probabilistic modeling of spatially distributed data. Spatial LDA is able to reduce complex assemblies of intermixed entities into distinct component communities (“topics”) while accounting for uncertainty and missing data; it has performed well on other multiplexed tissue imaging datasets^65, 66^. We separated CRC specimens in Cohort 1 into tumor and adjacent normal tissue using H&E data and an ML/AI model^51^ and then performed spatial LDA at the level of individual IF markers on cells in the tumor region (**Fig. 5a**). This yielded 12 spatial features (topics) that recurred across the dataset (the number of topics was optimized by calculating the perplexity; see Methods for details) (**Extended Data Fig. 7a**). Visual inspection of images by a pathologist confirmed that marker probabilities matched those computed for different topics and that the frequency distribution of each topic varied, sometimes substantially, among CRC samples (**Fig. 5b** and **Extended Data Fig. 7b**). The strongest correlations between topics and PFS for Cohort 1 were found to be -0.52 (p < 0.001) for Topic 7, comprising pan-cytokeratin and E-cadherin positivity (with little contribution from immune cells) and +0.57 (p < 0.001) for Topic 11, comprising CD20 positivity with minor contributions from CD3, CD4, and CD45 (**Fig. 5b-5f** and **Extended Data Fig. 7a**). In contrast, topics involving the proliferation marker Ki-67^+^ (Topic 6), PD-L1 positivity (Topic 9), or immune cells markers (CD45^+^ or CD45RO^+^; Topics 3 and 10) exhibited little or no correlation with progression-free survival (**Extended Data Fig. 7a**).

**Fig. 5.**
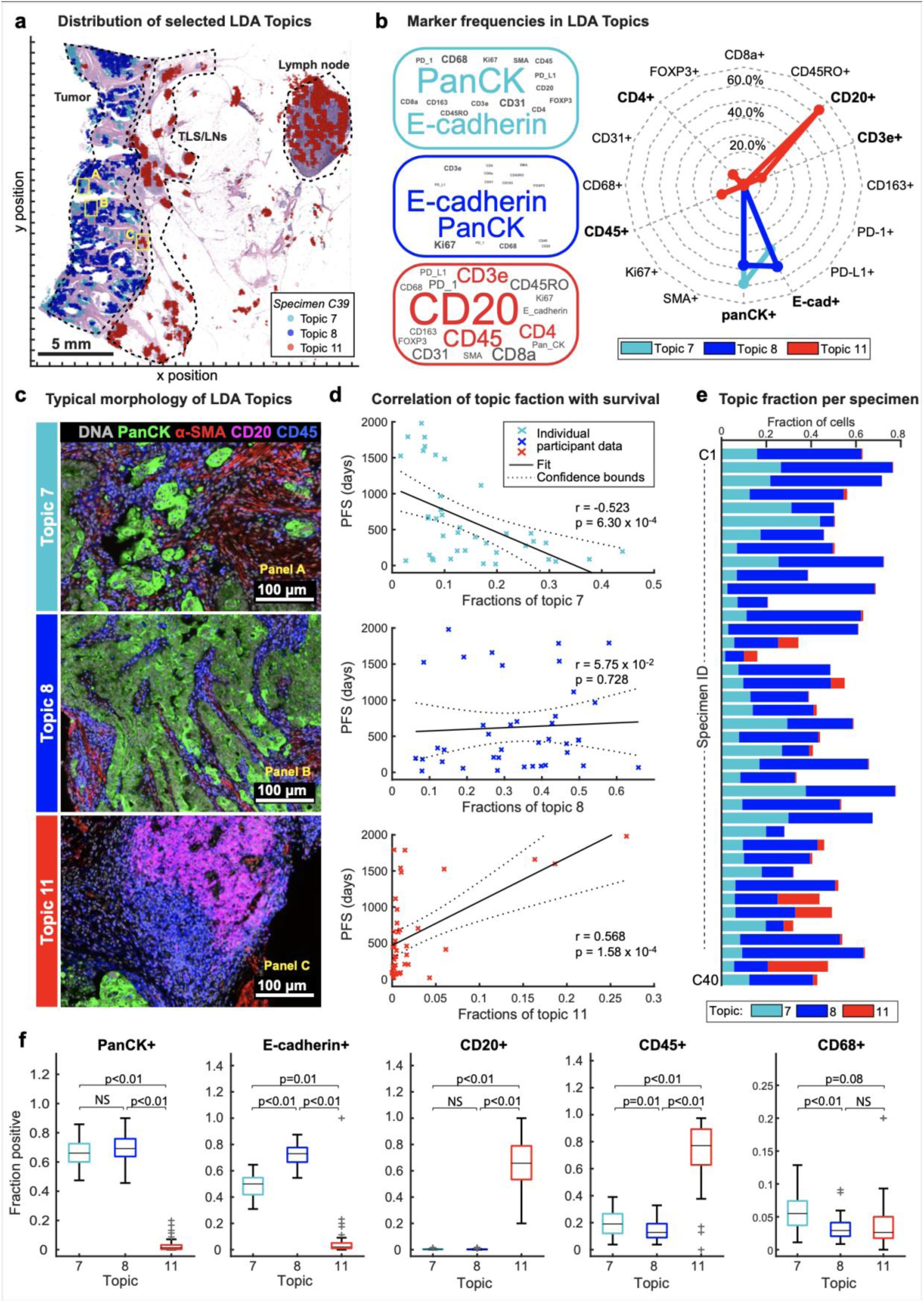
Bottom-up development of a tumor-intrinsic image feature model. **a,** Positions in specimen C39 of three selected topics identified using Latent Dirichlet Allocation (LDA). Topic locations are overlaid on an H&E image; Scalebar 5 mm. **b,** Left: Markers making up selected LDA topics as shown with size of the text proportional to the frequency of the marker but with colored text scaled by 50% for clarity; Radar plot indicating the fraction of cells positive for each marker in Topics 7, 8, and 11 (data for all others topics shown in **Extended Data Fig. 7**). **c,** Immunofluorescence images showing expression of pan-cytokeratin, α-SMA, CD20, and CD45 for the indicated LDA topics. The position of each image frame is denoted by the yellow boxes in panel a. Scalebars 100 µm. **d,** Pearson correlation plots of progression-free survival (PFS) and Fraction of Topic 7, 8 and 11 in 40 CRC patients. Topic 11 corresponded to TLS, whose presence is known to correlate with good outcome^87^. **e,** Fraction of Topics 7, 8, and 11 in colorectal cancer specimens C1-C40. **f,** Box-and- whisker plots showing fractions of Topic 7, 8, and 11 positive cells for indicated markers; midline = median, box limits = Q1 (25th percentile)/Q3 (75th percentile), whiskers = 1.5 inter-quartile range (IQR), dots = outliers (>1.5IQR)). Pairwise t-test p values indicated.

Given the correlation of Topic 7 with PFS, we constructed a Kaplan-Meier curve for tumors having a proportion of Topic 7 below the 50^th^ percentile versus those above this threshold (including all cells in the specimen). Imposing this threshold yielded model IFM3 which, on Cohort 1, yielded HR = 0.26 (**Fig. 6a**; CI 95%: 0.11 – 0.63; p = 2.98 × 10^-4^; note that the value of the threshold was not critical over the range of 50% – 75%) (**Fig. 6a** and **Extended Data Fig. 8a**). When we tested IFM3 on Cohort 2 we observed even better performance (HR = 0.07; CI 95%: 0.02 - 0.24; p = 5.6 × 10^-4^; **Fig. 6b**), suggesting that the model had not been over trained. We conclude that spatial-LDA had discovered – via unsupervised analysis of high-plex IF data – a tumor-intrinsic feature distinct from immune infiltration that was significantly associated with poor patient survival.

**Fig. 6.**
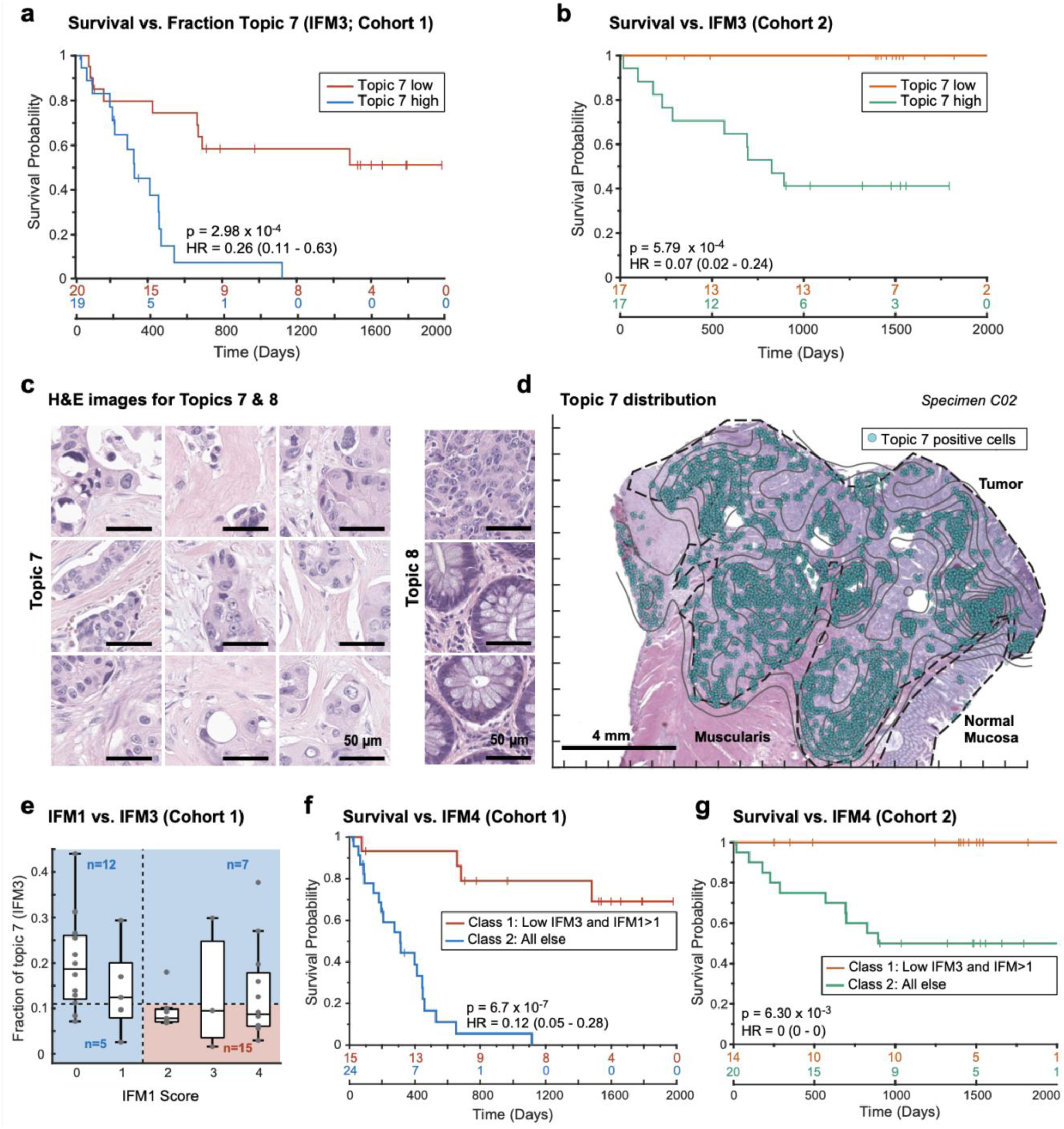
LDA Topic 7 corresponds to aggressive tumor regions and is correlated with poor outcomes. **a&b,** Kaplan Meier plots of PFS based on the fraction of Topic 7 present in the tumor domain and stratified as follows: high class: above median (50 percentile) of all cases, and low class: below median (HR, hazards ratio; 95% confidence interval; logrank p-value) for **a,** 40 CRC Cohort 1 patients and **b,** 34 CRC Cohort 2 patients. **c,** Representative images of Topic 7 (left) and Topic 8 (right) extracted from all specimens using a convolutional neural network (GoogLeNet) trained on LDA data. **d,** Spatial map of LDA Topic 7 and H&E image from colorectal cancer sample C02. **e,** Plot of fraction of Topic 7 (IFM3) versus IFM1 score for 40 CRC patients. **f&g,** Kaplan Meier plots stratified using IFM4 which was binarized as follows: class 1: IFM1 high and Topic 7 (IFM3) low group; class 2: all other patients – i.e., either low IFM1 and/or high Topic 7 (IFM3) (HR, hazards ratio; 95% confidence interval; logrank p-value), for **g,** Cohort 1(40 CRC patients) and **h,** Cohort 2 (34 CRC patients).

One limitation of this, and many other models built using ML methods such as spatial LDA, is poor interpretability. In the case of Topic 7, the primary molecular features were pan-cytokeratin and E-cadherin positivity, but Topic 8 was similar in composition while exhibiting no correlation with PFS (r = 0.01; **Fig. 5c, 5f** and **Extended Data Fig. 7a**). To identify the tumor histomorphology corresponding to these topics, we transferred labels from IF to the same section H&E images, trained a convolutional neural network (CNN) on the H&E data, and inspected the highest scoring tumor regions (**Extended Data Fig. 8b**). In the case of Topic 7, these were readily identifiable as regions of poorly differentiated/high-grade tumor with stromal invasion (**Fig. 6c** and **6d**). In contrast, Topic 8 consisted predominantly of intestinal mucosa with a largely normal morphology and some areas of well-differentiated tumor (**Fig. 6c** and **Extended Data Fig. 8c**). When we inspected Orion and CyCIF images of specimens with a high proportion of Topic 7 (e.g., patient C06, **Extended Data Fig. 9**) we found that the E-cadherin to pan-cytokeratin ratios were low relative to normal mucosa or Topic 8 (expression of Na,K-ATPase, another protein found on the plasma membranes of colonic epithelial cells, was also low). These are features of cells undergoing an epithelial-mesenchymal transition (EMT), which is associated in CRC with progression and metastasis^67^. However, follow-on CyCIF imaging showed that some features of EMT, such as low proliferation and increased expression of EMT-associated transcriptional regulators (e.g., ZEB1)^68^, were not generally observed in Topic 7-positive cells: the proliferation index was high (40-50% Ki67 and PCNA positivity) and staining for ZEB1 was low in tumor cells (even though ZEB1 was easily detected in nearby stromal cells with mesenchymal differentiation – compare yellow and white arrows; **Extended Data Fig 9**). Thus, even though the molecular and morphological features of Topic 7 were consistent with each other, H&E morphology was more readily interpretable with respect to long established features of CRC progression. It has been observed previously that interpretability increases confidence in a potential biomarker and substantially improves its chances of clinical translation^69^.

Only about one-third of patients in Cohort 1 scored high for IFM1 and low for IFM3 (the combination correlated with the longest PFS; **Fig. 6e**), so we reasoned that it would be effective to combine the two models. Using a composite model (IFM4), we observed excellent discrimination between progressing and non-progressing CRC patients with HR = 0.12; (95% CI = 0.05 to 0.28; p = 6.7 × 10^-7^) (**Fig. 6f**). Statistically significant results were also obtained from Cohort 2 using a model trained on Cohort 1 (**Fig. 6g**).This demonstrates that immunological and tumor-intrinsic features of cancers arising from top-down and bottom-up analysis can be effectively combined to generate prognostic models with high predictive value. Of note, no parameter tuning (e.g., setting thresholds for positivity) was involved in the generation of IFMs 1-3 or the highly performative IFM4 hybrid model, reducing the risk of over-training. Experience with Immunoscore shows that parameter tuning using larger cohorts of patients can further boost performance.

## DISCUSSION

In this paper, we describe an approach to multimodal tissue imaging that combines high-plex, subcellular resolution IF with conventional H&E imaging of the same cells and show that the approach can generate performative progression biomarkers of a common type of cancer in a whole-slide format suitable for clinical translation. The approach required developing a new Orion instrument, fluorophores, and protocols to enables both one cycle (single-shot) and two cycle high-plex IF data acquisition while preserving the sample for same-section H&E imaging. We show that such multimodal tissue imaging is reproducible across performance sites and has substantial benefits for human observers and machine-learned models. Most obviously, it facilitates the use of extensive historical knowledge about tissue microanatomy (derived from H&E images) for the interpretation of molecular data derived from multiplexed molecular imaging. We demonstrate this directly by showing that both human experts and ML algorithms can exploit H&E images to classify cell types and states that are not readily identifiable in multiplexed data given inevitable limitations in antibody variety. H&E and autofluorescence imaging are also effective at characterizing acellular structures that organize tissues at mesoscales (e.g., the elastic lamina of the vessel wall). At the same time, by overlaying molecular data on H&E images we show that it is possible to discriminate cell types that have similar morphologies but different functions. The ability of molecular data to label cell types in H&E images is expected to be advantageous in supervised learning for ML/AI modeling^7,70^ as well as the use of H&E data to analyze “black box” ML models trained on molecular data. The topic of black box versus interpretable AI is a major point of discussion in medicine in general^71^, but in the case of pathology it is highly likely that interpretability will improve uptake, facilitate further research, and improve generalizability across cohorts.

The Orion described here instrument currently supports up to 20-plex data simultaneous data acquisition (including DNA and one or more autofluorescence channels), but we have found that 18-plex data collection is more robust – hence its use in this paper (see Materials and Methods for a detailed discussion of this point). It is nonetheless likely that several additional channels can be added to the approach as we identify fluorophores more optimally matched to available lasers and optical elements. We show successful 18-plex Orion imaging of 30 types of cancer, diseased tissues, and normal tissues available as TMA cores or whole slide specimens, demonstrating that the Orion method is widely applicable. Of course, the combination of antibodies in our colorectal cancer panel is not optimal for such a wide variety of tissues, but substitution of a few antibodies is expected to yield near-optimal panels for many cancers of epithelial origin. Moreover, a wide range of commercial antibodies developed for IHC and IF imaging of tissues are suitable for conjugation with ArgoFluors and use in Orion panels; the only practical limitation to development of these panels is the time needed to test conjugated antibodies is various combinations and then validate panel performance and stability.

We show that it is possible to perform cyclic data acquisition using the Orion approach as well as Orion followed by CyCIF, thereby increasing the number of molecular channels dramatically. Cyclic Orion is particularly well suited to discovery research in which 20-40 plex imaging is increasingly common^72^. However, H&E staining must be performed after all IF is complete, and we find that H&E image quality goes down as IF data acquisition extends beyond 2-4 cycles (although additional protocol optimization may extend this). For diagnostic applications, the imperative for simplicity and reliability is greater than in a research setting, and our data suggest that performative image-based prognostic tests may require only a subset of the channels available to Orion (speculatively 8-14 channels) with attendant reductions in test complexity and cost.

### Complementarity of same-section Immunofluorescence and H&E imaging

It is not surprising that multiplexed molecular data from IF images add information to H&E imaging. More surprising are the many cell types and structures that are difficult to identify in multiplexed images and readily identified in H&E images by histopathologists or the ML algorithms they train. This includes acellular structures, cell types for which good markers are not readily available, highly elongated and multi-nucleated cells that are difficult to segment with existing algorithms (e.g., muscle cells), and – most remarkably – tumor cells themselves. Many tumor types lack a definitive cell-type marker, and even when such markers are available, some cells in a tumor are observed to express these markers poorly or not at all, likely due to sub-clonal heterogeneity^73^. In contrast, pathologists are skilled at identifying dysplastic and transformed cells in H&E images. Thus, H&E imaging in combination with ML models is potentially more reliable than IF imaging using molecular markers for the identification of some types of tumor cells. Conversely, many immune cell types cannot be reliably differentiated using H&E images, and their presence can also be difficult to discern when cells are crowded; the use of IF lineage markers provide critical new information in these cases.

The complementary strengths of H&E and IF imaging can be exploited by ML/AI algorithms that are increasingly used to process tissue images in clinical and research settings^70^. For example, we show that models trained to recognize disease-associated structures in H&E images, which is an area of intensive development in digital pathology^74^, can improve the analysis and interpretation of multiplexed IF data. The converse is also true: IF images can be used to automatically label structures in H&E images (e.g., immune cell types) to assist in supervised learning on these images. This is a significant development because the labor associated with labeling of images – currently by human experts – is a major barrier to the development of better ML models. Combined H&E and IF images will be of immediate use in ML-assisted human-in-the loop environments that represent the state of the art in image interpretation in a research setting^75^.

### Using the Orion approach to advance prognostic and predictive biomarkers

A surprising number of pathology workflows involve staining serial sections of a specimen each with one IHC biomarker on followed by manual inspection of images by histopathology experts. The Orion approach used in conjunction with open-source software pipelines^41^ has the potential to automate these workflows and also provide new molecular insight into tumor features already known from H&E data to be prognostic of tumor progression^76^. For example, Immunoscore is a pathology-driven (top-down) clinical test that uses H&E and IHC data to determine the distribution of specific immune cell types at the tumor margin and predict outcome (time to recurrence) for patients with CRC. In this paper, we reproduced the logic of Immunoscore and used automated scripts to show that it is possible to improve upon it using additional immune markers and a single round of data acquisition (as measured by Hazard Ratios computed from PFS data; see limitations section below)^77^.

In a distinct but complementary bottom-up approach, we use spatially sensitive statistical model (LDA) of IF data to identify cell neighborhoods significantly associated with CRC progression. The top-performing feature in this case is tumor-cell intrinsic and is based on the distributions of cytokeratin and E-cadherin, two epithelial cell markers. Precisely why this is feature is prognostic is unclear from IF data alone: other features involving similar markers are not predictive. However, inspection of corresponding H&E data (and training of an ML model) showed that LDA had identified local tumor morphologies typical of poorly differentiated/high-grade tumor with stromal invasion, increasing our confidence in the model. Because the features in the tumor-intrinsic model were distinct from and uncorrelated with the immune markers in Immunoscore, combining the two sets of features significantly improved the hazard ratio relative to either model used alone. We therefore anticipate that many opportunities will emerge for joint learning from H&E and IF data using adversarial, reinforcement, and other types of ML/AI modeling for research purposes, development of novel biomarkers, and analysis of clinical H&E data at scale^6^. The immediate availability of Orion as a commercial platform and our use of open-source software and OME (Open Microscopy Environment)^78^ and MITI (Minimum Information about Tissue Imaging)^79^ compliant data standards makes the approach we describe readily available to other investigators.

### Limitations

Although the images in this paper represent the largest dataset collected to date using high-plex whole-slide IF imaging, the number of specimens and the composition of the cohort is insufficient for IFMs to be considered validated biomarkers or clinical tests^80^. Systematic meta-analysis has identified a range of factors that negatively impact the reliability and value of prognostic biomarkers^81^, particularly those based on new technology and multiplexed assays^82^. In the current work, specific limitations include a relatively small cohort size, the absence of pre-registeration^83^, and the acquisition of specimens from a single institution. The limited number of specimens in the current study, as compared to conventional practice in histopathology (in which study of 500 cases is not uncommon), makes it impossible to fully control for all relevant covariates (e.g., depth of invasion, sex, age, race, clinical stage etc.). Moreover, to enable better detection of image features associated with progression, more progressors were included in our cohort than would be observed in an unselected population, biasing the cohort to more serious disease (the two-year disease-free survival for Stage III colon cancer in a 12,834 patient multi-center cohort was reported to be ∼80%^84^ but it is only ∼50% in our cohort). These and other concerns will be addressable as we gain access to larger and more diverse collections of tissue blocks from which fresh sections can be cut and multi-modal imaging performed. With all of the advantages attendant to automated data acquisition and ML-based image analysis we anticipate that it will be feasible to progress in a few years to validated clinical tests that can be added to colorectal cancer treatment guidelines^60^, substantially improving opportunities for personalized therapy.

## Supporting information

Extended_figures_with_legends

ExtData_table_1

ExtData_table_2

ExtData_table_3

ExtData_table_4

ExtData_table_5

## ACKNOWLEDGEMENTS

This work was supported by NCI grants U54-CA225088 and U2C-CA233262 (PKS, SS), an NCI SBIR small business grant to RareCyte and PKS (R41-CA224503), and commercial investment from RareCyte; image processing software and data science methods were developed with support from the Bill and Melinda Gates Foundation grant INV-027106, a Team Science Grant from the Gray Foundation, David Liposarcoma Research Initiative, Emerson Collective, and Ludwig Cancer Research. SS is supported by the BWH President’s Scholars Award. We are grateful to all members of the HMS Laboratory of Systems Pharmacology (LSP) engaged in tissue imaging (see https://www.tissue-atlas.org/), to Joe Victor, and to members of the RareCyte software and hardware development teams.

## AUTHOR CONTRIBUTIONS

J.R.L, Y.C., D.C., J.C., and E.M performed experiments and imaging. J.R.L., Y.C., D.C., J.C., S.C., C.Y., S.R., and T.G. performed data analysis. P.K.S., S.S., T.G., J.R.L., Y.C., and J.B.T. wrote the paper and all authors reviewed drafts and the final manuscript. J.B.T., J.R.L., and Y.C. prepared the figures. K.L.L., S.J.R., and S.S. supervised clinical research, and S.R., T.G., S.S., and P.K.S. supervised the overall research.

## COMPETING INTERESTS

PKS is a co-founder and member of the BOD of Glencoe Software, a member of the BOD for Applied Biomath, and a member of the SAB for RareCyte, NanoString, and Montai Health; he holds equity in Glencoe, Applied Biomath, and RareCyte. PKS is a consultant for Merck and the Sorger lab has received research funding from Novartis and Merck in the past five years. YC is a consultant for RareCyte. DC, JC, EM, SR, and TG are employees of RareCyte. The DFCI receives funding for KLL’s research from the following entities: Amgen, Travera, and X4. DFCI and KLL have patents related to molecular diagnostics of cancer. SJR receives research support from Bristol-Myers-Squibb and KITE/Gilead. SJR is on the Scientific Advisory Board for Immunitas Therapeutics. The other authors declare no outside interests.

## MATERIALS AND METHODS

### Ethics and tissue cohort

Our research complies with all relevant ethical regulations and was reviewed and approved by the Institutional Review Boards (IRB) at Brigham and Women’s Hospital (BWH), Harvard Medical School (HMS), and Dana Farber Cancer Institute (DFCI). Formaldehyde-fixed and paraffin-embedded (FFPE) tissue samples were used after diagnosis and informed written patient consent under Dana-Farber Cancer Institute IRB protocol 17-000. The study is compliant with all relevant ethical regulations regarding research involving human tissue specimens. Two cohorts from same biobank were assembled, the first with 40 patients with state II-IV CRC, then the second with 34 patients. Samples were collected at the time of initial diagnosis.

### Tissue preparation

Blocks of FFPE tonsil (AMSBIO, cat# 6022CS) and lung adenocarcinoma (AMSBIO, cat# 28004) and colorectal adenocarcinoma from the BWH Pathology Department archives were cut at 5 µm thickness using a rotary microtome and the sections were mounted onto Superfrost™ Plus microscope glass slides (Thermo Fisher, Catalog No.12-550-15). The slides were dried at 37°C overnight and baked at 59°C for one hour. Slides were stored at 4°C until use.

### Fluorophores for Orion™ imaging

The Orion™ instrument is designed to work with an optimized set of fluorophores from RareCyte, branded as ArgoFluor™ dyes whose emission peaks cover the spectrum from green to far-red (**Extended Data Table 2**). Although the instrument can also be used with other commercially available dyes, the ArgoFluor™ dyes have been strategically chosen based on a combination of properties that include resistance to photobleaching, narrow excitation and emission spectra, and high quantum efficiency. To date, the company has optimized 18 ArgoFluor dyes, with others in development.

### Immunofluorescence antibodies

Antibodies were obtained in carrier-free PBS and conjugated directly to either biotin for α-SMA, digoxygenin for pan-cytokeratin or to ArgoFluor™ dyes (RareCyte, Inc.) using amine conjugation chemistry. After determining labeling efficiency using absorbance spectroscopy, the conjugated antibodies were diluted in PBS-Antibody Stabilizer (CANDOR Bioscience GmbH, Catalog No. 130050) to a concentration of 200 µg/mL. Antibodies used in immunofluorescence studies are listed in the **Extended Data Table 2**.

### Immunofluorescence staining

Slides were de-paraffinized and subjected to antigen retrieval for 5 minutes at 95°C followed by 5 minutes at 107°C, using pH8.5 EZ-AR 2 Elegance buffer (BioGenex, Catalog No. HK547-XAK). To reduce tissue autofluorescence, slides were placed in a transparent reservoir containing 4.5% H_2_O_2_ and 24 mM NaOH in PBS and illuminated with white light for 60 minutes followed by 365 nm light for 30 minutes at room temperature as previously described^15^. Slides were rinsed with surfactant wash buffer (0.025% Triton X-100 in PBS), placed in a humidified stain tray, and incubated in Image-iT™ FX Signal Enhancer (Thermo Fisher, Catalog No. I36933) for 15 minutes at room temperature. After rinsing with surfactant wash buffer, the slides were placed in a humidity tray and stained with the panel of fluor- and hapten-labeled primary antibodies in PBS-Antibody Stabilizer (CANDOR Bioscience GmbH, Catalog No.130 050) containing 5% mouse serum and 5% rabbit serum for 2 hours at room temperature. Slides were then rinsed again with surfactant wash buffer and placed in a humidified stain tray and incubated with Hoechst 33342 (Thermo Fisher Catalog no. H3570), ArgoFluor™ 845 mouse-anti-DIG, and ArgoFluor™ 875-conjugated streptavidin in PBS-Antibody Stabilizer containing 10% goat serum for 30 minutes at room temperature. The slides were then rinsed a final time with surfactant wash buffer and PBS, coverslipped with ArgoFluor™ Mounting Media (RareCyte, Inc.) and dried overnight.

### ArgoFluor™-antibody conjugate stability testing

Antibody accelerated-aging studies were performed to determine ArgoFluor™-antibody conjugation stability. Reagent stability was measured using the ratio of quantitative metrics obtained with the accelerated condition (21.6°C) to those obtained with the storage condition (-20°C). Tissue validation (Orion IF): Single-cell mean fluorescence intensity (MFI) data obtained by imaging FFPE tonsil stained with the ArgoFluor™ conjugate was gated using a Gaussian mixture model to obtain the percent of positive cells and S:B values (S and B refer to the MFI of cells with values above (S, Signal) and below (B, Background) the gated threshold). Fluor stability (Orion IF): Single bead MFI data was obtained by imaging Ig-capture beads incubated with (S) or without (B) the ArgoFluor™ conjugate. Binding stability (Flow Cytometry): Intensity data from peripheral blood mononuclear cells (PBMC) stained with the ArgoFluor conjugated antibody was manually gated to obtain % Positive and S:B values (S and B refer to the MFI of cells with values above (S) and below (B) the gated threshold).

### The Orion method and instrumentation

The Orion instrument was designed with the following performance goals: (1) whole-slide imaging; (2) rapid single-pass data collection; (3) sub-cellular imaging resolution; (4) sufficient immunoprofiling depth; (5) bright-field imaging; (6) optical and mechanical stability for accurate image tile stitching; and (7) compatibility with established image data standards and formats. ArgoFluor™-conjugated antibodies along with Hoechst dye and tissue autofluorescence were excited by seven laser lines, ranging from 405 to 730 nm (**Extended Data Table 2**). To separate the overlapping emission spectra, images were captured through a set of nine bandpass filters, which can achieve a tunable narrow band detection window (10 - 15 nm) throughout the spectrum from 425 nm to 894 nm. For a specific sample, the detection bands were chosen to optimize color separation, implemented with RareCyte Inc.’s Artemis™ software. Tuning of these filters is based on the well-known fact that the spectrum of a thin-film interference filter shifts toward shorter wavelengths when the angle of incidence shifts away from 0 degrees (orthogonal to the filter surface). The filters were motorized such that any narrow band of 10 - 15 nm can be achieved across the entire fluorescence spectrum. Narrow bandpass emission channels improve specificity; the resulting lower signal is overcome by using high power excitation lasers, which yield power at the sample plane ranging from 270 to 600 mW, more than 10 times greater than a typical fluorescence microscope.

### Considerations in the development of Orion antibody panels

High-plex imaging exploits the fact that the greater the number of features collected, the greater the ability to distinguish lineages and states at a single-cell level. The ability of the Orion imaging platform to discriminate among multiple antibody-fluorophore conjugates is dependent on the degree of spectral overlap among the fluorophores, the intensity and spectral profile of overlapping autofluorescence or background signals, and the difference between the most intense staining of highly expressed proteins and the weakest stain of low abundance proteins. Panel design with the Orion platform involves assigning biomarkers to channels with the appropriate sensitivity ranges while managing spectral overlap between markers that are co-localized. Orion imaging technology is compatible with 20-plex one-shot fluorescence image acquisition (19 antibodies plus Hoechst nuclear stain) and the necessary research into ArgoFluor is ongoing to achieve this on a routine basis. In the current work we found that 17-plex panels were easier to achieve at an acceptable SNR given the properties of tonsil and CRC tissue. We anticipate that, with relatively few additions and substitutions, the panel we developed for CRC will work well other common tumor types (e.g., lung, breast, melanoma). In cases in which more precise immunophenotyping is desired, a second cycle Orion panel of similar complexity is possible. However, it is important to note that both autofluorescence imaging and the use of ML on H&E images have the potential to generate data additional data on cell types and states – potentially equivalent to 10 or more antibody channels. The prognostic image feature models we describe in this paper could also be acquired using as few as 8-12 channels. Thus, optimal Orion imaging and staining strategies in both a research and clinical setting are likely to rely on the use of both pre-set, high-plex, more difficult to optimize panels and lower-plex, lower-cost, “mix and match” panels.

### One-shot antibody IF imaging with the Orion instrument

Whole slides were scanned using the Orion instrument using acquisition settings optimized for the specific antibody panels. Briefly, acquisition channel parameters were defined for each biomarker plus an additional channel dedicated to tissue autofluorescence, and included excitation laser, emission center wavelength (CWL), and exposure times. The nuclear channel was scanned at low resolution to identify tissue boundaries, followed by surface mapping at 20X to find the tissue in the z-axis. Whole tissue was acquired at 20X following the surface map within the specified tissue boundaries by collecting all channels for a single field of view (FOV) before proceeding to the next partially overlapping FOV. Raw image files were processed to correct for system aberrations, then signal from individual targets were isolated to separate channels using the Spectral Matrix obtained with control samples, followed by stitching of FOVs to generate a continuous open microscopy environment (OME) pyramid TIFF image.

### Same Section H&E staining and imaging

After Orion imaging was complete, slides were de-coverslipped by immersion in 1x PBS at 37°C until the coverslips fell away from the slide. Slides were rinsed in distilled water for 2 minutes, then stained by a routine regressive H&E protocol using Harris Hematoxylin (Leica, Catalog No. 3801575) and alcoholic eosin Y (Epredia, Catalog No. 71211). Coverslipping was performed with toluene-based mounting media (Thermo Scientific, Catalog No. 4112). After drying for 24 hours, slides were scanned on an Orion system in brightfield mode, using the same scan area used for IF image acquisition. H&E images were also acquired using an Aperio GT450 microscope (Leica Biosystems), and the H&E images were registered to the IF images using ASHLAR^48^ and PALOM software (https://github.com/Yu-AnChen/palom).

### Pathology annotation of H&E images performed after Orion immunofluorescence imaging

H&E images were annotated by a board-certified anatomic pathologist (SC and SS). The histologic features of each tissue section were defined and labeled in OMERO PathViewer software on whole slide images according to morphologic criteria^85^ including normal mucosa, hyperplastic mucosa, adenomatous mucosa (tubular or serrated), invasive adenocarcinoma (tumor), lymphovascular invasion (LVI), peri-neural invasion (PNI), secondary lymphoid structures/Peyer’s patches (SLS), tertiary lymphoid structures (TLS), lymphoid aggregates (without identifiable germinal center formation), lymph nodes. Tertiary lymphoid structures were morphologically defined by the presence of a lymphoid aggregate with germinal center formation and an anatomic distribution and appearance inconsistent with a secondary lymphoid structure (Peyer’s patch or lymph node).

### CyCIF imaging

Tissue-based cyclic immunofluorescence (CyCIF) was performed as previously described^15^ following the methods available in protocols.io (<u>dx.doi.org/10.17504/protocols.io.bjiukkew</u>). Data from specimens C1-C17 was acquired as previously reported^22^ and computed cell counts were compared in this study with cell counts derived from Orion images of adjacent sections from the same specimens. A BOND RX Automated Slide Stainer was used to bake FFPE slides at 60°C for 30 minutes. Dewaxing was performed using Bond Dewax solution at 72°C, and antigen retrieval was performed using BOND Epitope Retrieval Solution 1 (Leica Biosystems) at 100°C for 20 minutes. Slides then underwent multiple cycles of antibody incubation, imaging, and fluorophore inactivation to perform the CyCIF process. All antibodies were incubated overnight at 4°C in the dark. Slides were stained with Hoechst 33342 for 10 minutes at room temperature in the dark following antibody incubation in every cycle. Coverslips were wet-mounted using 200µL of 10% Glycerol in PBS prior to imaging. Images were taken using a 20X objective (0.75 NA) on a CyteFinder™ slide scanning fluorescence instrument (RareCyte Inc. Seattle WA). Fluorophores were inactivated by incubating slides in a 4.5% H_2_O_2_, 24mM NaOH in PBS solution and placing under an LED light source for 1 hr. For CyCIF after Orion imaging, slides were immersed in 1x PBS at 37°C until the coverslips fell away from the slide. The standard CyCIF method was subsequently performed on these slides.

### Immunohistochemistry

FFPE sections were de-paraffinized, dehydrated, and endogenous peroxidase activity was blocked. Antigen retrieval was performed for 20 minutes at 100°C, at pH9, using BOND Epitope Retrieval Solution 2 (Leica Biosystems). Detection was achieved using a Bond Polymer Refine Detection® DAB chromogen kit and counterstained with hematoxylin. Slides were scanned using a RareCyte CyteFinder instrument. Primary antibodies used in immunohistochemistry are listed in **Extended Data Table 2**.

### Orion image processing data quantification

*Image stitching and segmentation.* Image data processing was performed using MCMICRO modules^41^. Briefly, stitched, registered, illumination and geometric distortion corrected images were generated by the Orion platform. Single-cell segmentation was performed with UNMICST2 and cell masks were generated by 5-pixel dilation of the nucleus masks. Mean intensity of each channel and morphological features were quantified for each cell masks. Image and data analysis was performed using customized scripts in Python, ImageJ and MATLAB. All code is available on GitHub (https://github.com/labsyspharm/orion-crc).

### Analysis of channel crosstalk

#### Single-plex tonsil images

Tonsil FFPE sections stained with single antibody-ArgoFluor underwent standard acquisition and extraction process using the Orion instrument. The pixel intensities of all 18 channels from 17 samples were used to quantify bleed through of a given antibody-ArgoFluor complex to the other channels before and after spectral extraction.

#### 18-plex tonsil image

Pearson’s correlation coefficients between all channel pairs were computed using pixel intensities in the 18-plex tonsil image before and after spectral extraction.

### Computational analysis of Orion images and derivation of image feature models

#### IFM computation from Orion data

IFM1 was designed to replicate the logic of the Immunoscore method and was calculated in a semi-automated manner using Orion data. In brief, quantitative data of tumor and immune markers (pan-cytokeratin, CD3e, and CD8a) were gated for high and low cells using a Gaussian Mixture Model (GMM) and confirmed by inspection. After gating, the pan-cytokeratin^+^ cells were then used to generate tumor masks using a K-Nearest Neighbor (KNN) model (kernel size = 25 cells). The tumor margins were derived from tumor masks by expanding 100 microns in either direction from the point of stroma-tumor contact. The CD3^+^ and CD8^+^ fraction, defined as marker positive cells divided by the total of all successfully segmented cells of all types in either the tumor center (TC) or invasive margin (IM). Tumor and margins were enumerated independently in each sample. The median values of all samples were used as a cutoff to defined a subscore as follows: below the median scored as 0 and above the median scored as 1. The final IFM1 value was calculated by adding together the subscores for CD3 and CD8 positive cells in the TC and IM regions (see **Fig. 4b** for a flow diagram). The IFM1 score therefore ranged from 0 (CD3^+^ and CD8^+^ low in both regions) to 4 (CD3^+^ and CD8^+^ high in both regions). Similar logic was used to generated other combinations of IFMs. 13 selected immune markers (CD3, CD8, CD45, CD45RO, CD68, CD163, CD4, CD20, α-SMA, FOXP3, PD-1, PD-L1) were gated as described above, and 26 parameters (each marker in the tumor or tumor/stromal interface regions) were generated. The complete combination of 4 out 26 parameters was tested against PFS days for Hazard Ratio (HR). IFM2 was the 3^rd^ best IFM among those combinations, excluding the 1^st^ and 2^nd^ best combinations which had some of the same markers as IFM1 (i.e., CD3 and CD8); the difference in performance between the top performing models was insignificant.

#### Leave-one-out (LOO) test and bootstrapping analysis for IFM2

In the LOO test, the ranks of IFM1 and IFM2 were recalculated with the 40 set of samples (n = 39); each set left out one sample from the original cohort. The collections of ranks from IFM1 and IFM2 were then tested with pairwise t-test. For bootstrapping, the 500 set of randomly selected samples were used to recalculate the hazard ratios of IFM1 and IFM2 as described above. The collections of hazard ratios from IFM1 and IFM2 were then tested with the pairwise t-test. To adjust for multiple hypotheses, the Benjamini-Hochberg Procedure was used with FDR = 0.1.

#### Latent Dirichlet Allocation for IFM3 and IFM4

Latent Dirichlet Allocation (LDA) was used to compute spatial neighborhoods as described^22^. First, each sample was divided into “grids” of 200 microns^2^, and marker frequency was calculated in each grid. The summarized probabilities of all samples were then used to generate the LDA model with 12 topics using collapsed Gibbs sampling in MATLAB. The optimal topic number was determined via varying numbers (between 8 to 16) of topics and evaluating the goodness-of-fit by calculating the perplexity of a held-out set. After fitting a global LDA model, the individual samples were then applied with the same models to assign topics at the single-cell level.

### Convolutional Neural Network to identify IFM3 in H&E images

A publicly available DenseNet161 model (https://doi.org/10.1101/2021.12.23.474029) trained with the 100K CRC H&E dataset (https://doi.org/10.5281/zenodo.1214456) was used to classify the post-Orion H&E image patches (112 µm^2^) for all the CRC samples. WSI patch prediction was performed with TIAToolbox v1.1.0 (https://doi.org/10.1101/2021.12.23.474029) on a Windows PC with Nvidia GeForce GTX 1080 graphics card and using batch size = 32. Model performance was reported as F_1_ = 0.992. As described in the training dataset, there are 9 output classes: adipose (ADI), background (BACK), debris (DEB), lymphocytes (LYM), mucus (MUC), smooth muscle (MUS), normal colon mucosa (NORM), cancer-associated stroma (STR), colorectal adenocarcinoma epithelium (TUM).Scripts for reproducing the inference results can be found at https://github.com/labsyspharm/orion-crc). The transfer learning of a GoogLeNet model was done as follows. First, the patch images of 224 × 224 pixels^2^ were generated from post-Orion H&E images, and the LDA topics were assigned to each patch using Orion data. To exclude low confidence training data, only patches with more than 20 cells and the percentage of the dominant topic over 60% were used. The selected patches were than separated into training, validation, and test sets as the ratio 0.6:0.2:0.2. The training was done with MATLAB (version 2019b) and the results are shown in **Extended Data Fig. 8b**. Scripts and training data are available at https://github.com/labsyspharm/orion-crc. Training parameters are listed at **Extended Data Table 5**.

### Outcome analysis

For all survival analyses, we used a combined survival endpoint of progression-free survival that encompasses both time to disease recurrence for patients who underwent curative-intent resections (disease-free survival; PFS) and time to progression for patients with measurable disease (progression-free survival; PFS); we used PFS in this paper because it is more familiar. Outcome analysis was performed using Kaplan-Meyer estimation and log-rank test utilizing the MatSurv function in MATLAB^86^. Cutoffs for IFM1, IFM2, and IFM3 were selected at the median value of the entire cohort, and cutoff for IFM4 were selected based on IFM1 & IFM3 as described. Hazard ratios and confidence intervals were calculated with the log-rank approach: HR = (Oa/Ea)/(Ob/Eb), where Oa & Ob are the observed events in each group and Ea & Eb are the number of expected events^78^.

## DATA AVAILABILITY (AT PUBLICATION – SEE INFORMATION FOR REVIEWERS ABOVE)

Data used in the preparation of this manuscript are detailed in the Source Data file provided with the manuscript. All image and derived data are available without restriction via the NCI Human Tumor Atlas Network (HTAN) Portal (https://humantumoratlas.org/explore) in accordance with NCI Moonshot Policies. HTAN participant ID is listed in **Extended Data** Table 3. Access to processed and unprocessed data are available via an index page on GitHub that has been archived on Zenodo (https://zenodo.org/) - https://doi.org/10.5281/zenodo.7637655.

## CODE AVAILABILITY

All code is available under an MIT open-source license via an index page on GitHub that has been archived on Zenodo (https://zenodo.org/) - https://doi.org/10.5281/zenodo.7637655.

## Notes

https://labsyspharm.github.io/orion-crc/

### Summary of Updates

We have collected a substantial amount of new data and made extensive changes to the text to address the reviewers' concerns. Most importantly, we have collected data from 34 additional human CRC specimens so that a classic split could be performed between training and test data. I am happy to report that the image feature models (biomarkers) we describe performed as well on the new test data as on the training data, substantially strengthening the manuscript. We also demonstrate a cyclic approach to Orion data collection, increasing the number of molecular markers from 18 to 32, and address a range of other technical concerns raised by the reviewers.

