## Extended_figures_with_legends for "Multi-modal digital pathology for colorectal cancer diagnosis by high-plex immunofluorescence imaging and traditional histology of the same tissue section"

#### Extended Data Figure 1

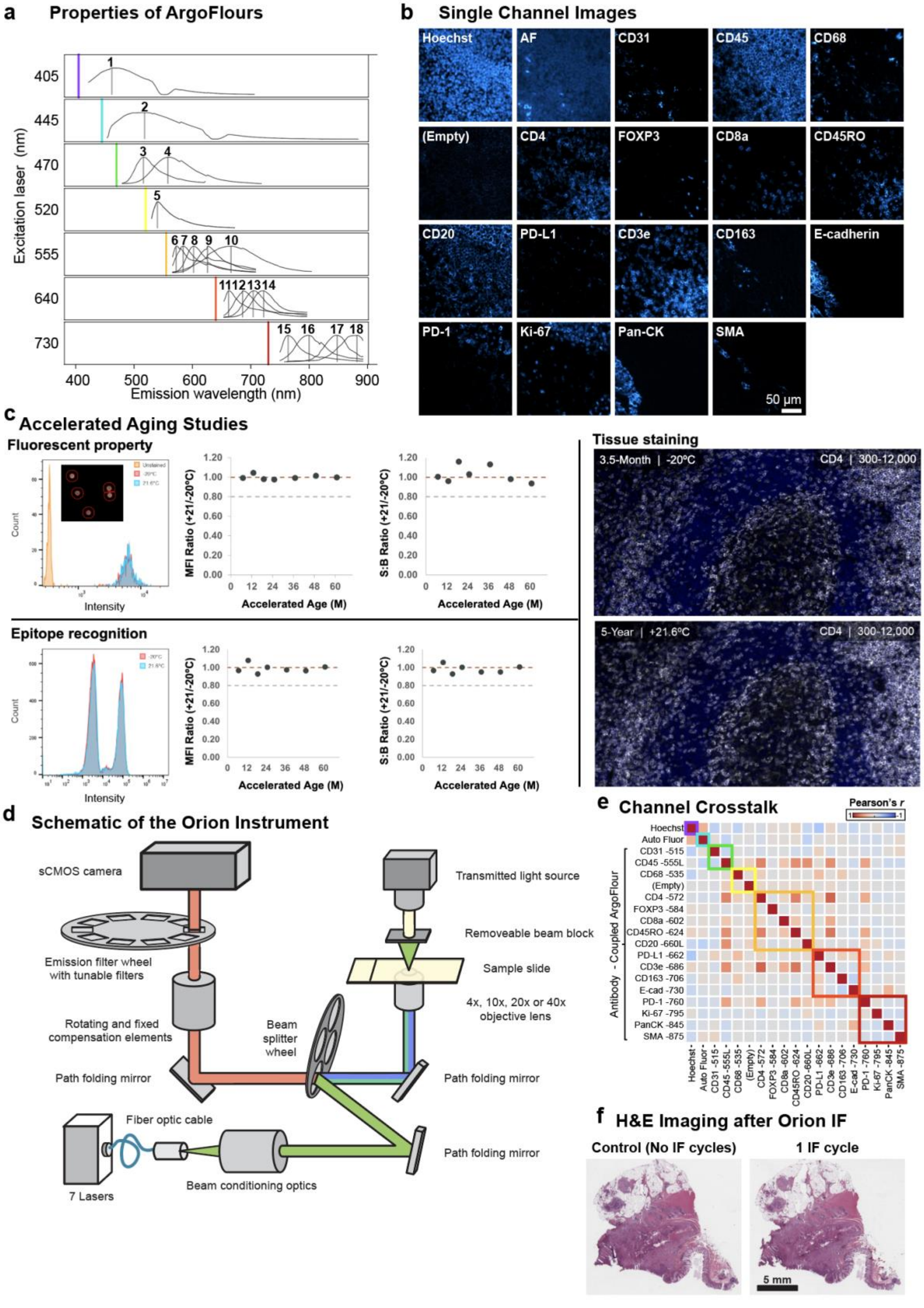


1108

**Extended Data Fig. 1 | Features of the fluorophores, signal extraction, antibodies, and instrumentation used in the Orion™ Method.**

**a,** Emission spectra of the ArgoFluor dyes used in this study with overlaid filter profiles. Each row shows fluorophores excited using the same laser (denoted by the colored vertical line). From left to right within each laser row: 405 laser (Hoechst 33342); 445 laser (Autofluorescence); 470 laser (ArgoFluor 515, ArgoFluor 555L); 520 laser (ArgoFluor 535, ArgoFluor 550); 555 laser (ArgoFluor 572, ArgoFluor 584, ArgoFluor 602, ArgoFluor 624, ArgoFluor 660L); 640 laser (ArgoFluor 662, ArgoFluor 686, ArgoFluor 706, ArgoFluor 730); 730 laser (ArgoFluor 760, ArgoFluor 795, ArgoFluor 845, ArgoFluor 875). For the 405-laser data collection, tonsil tissue stained with Hoechst 33342 was used as the sample. For the 445-laser data collection, unstained lung tissue was used. Single color Ig-capture beads generated by incubation with antibodies conjugated to the indicated ArgoFluor dye were used as the sample for all other collections. For each sample, data was collected into multiple Orion channels spanning a wide range of wavelengths (in 2 nm center wavelength increments). **b,** Channel images of FFPE tonsil section stained, imaged, and processed with Orion platform showing distinct spatial patterns with minimal channel crosstalk. **c,** Stability of fluorophore and of epitope recognition in solution and in tissues is shown for ArgoFluor 572 conjugated anti-CD4 antibody. Quantitative stability metrics were generated from three different assays to compare reagents stored at an accelerated aging condition (+21.6ºC) to reagents stored at the recommended condition (-20ºC) based on the Arrhenius equation (storage for 3.5 months at the accelerated aging condition is equivalent to 5 years at the recommended storage condition). *Fluorochrome stability*: The intensity of Ig-capture beads incubated with (signal) or without (background) antibody was measured from images scanned with the Orion system. The histogram overlay shows the intensity distribution for unlabeled beads (orange) and for beads incubated with antibody stored for 3.5 months at -20ºC (red) or +21.6ºC (blue). The mean fluorescence intensity (MFI) was obtained from these distributions, as well as the MFI signal-to-background (S:B) ratios. The dot plots show accelerated-to-real time CD4 MFI ratios (left plot) and S:B ratios (right plot) across 7 computed time points. *Antibody binding stability*: Human peripheral blood mononuclear cells (PBMC) were stained with accelerated-aged (blue) or real-time-aged (red) ArgoFluor 572 conjugated anti-CD4 antibody and analyzed using flow cytometry (3.5-month real-time / 5-year accelerated time point shown in histogram. The MFI was obtained for the positive (signal) and negative (background) populations, allowing derivation of S:B ratios. The dot plots show accelerated-to-real time CD4 MFI ratios (left) and S:B ratios (right) across 7 time points. For tissue-based antibody stability testing, images of serial sections from FFPE tonsil stained with real-time aged (top)z and accelerated-aged (bottom) antibodies were obtained using the Orion system. Single cell segmentation and intensity measurements were obtained with QuPath software, and a Gaussian mixture model threshold was applied to determine positive cells (signal) from negative cells (background) to determine S:B for both conditions. These methods demonstrate equivalent performance for both storage conditions in the three assays (S:B ratio of 0.93 for the 3.5-month real-time / 5-year accelerated time point). **d,** Schematic of the Orion optical system. The Orion imaging system has fluorescence and brightfield imaging modes. *Fluorescence imaging*: The Orion system is a class 1 LASER product which uses 7-color LASER illumination (one at a time) to illuminate a sample on a microscope slide. The illumination beam emanates from the source in a fiber optic cable, then shaped with beam conditioning optics, and redirected via a beam splitter and path folding mirrors through an objective lens which focuses it onto the sample. Excitation light passing through the sample is stopped by a beam block preventing damage to the transmitted light source. Laser-excited fluorophores in the sample emit light that is collected by the objective lens. This light is redirected via the beam splitter and path folding mirrors through a tube lens for focusing, fixed and rotatable compensation elements for optical corrections, and a tunable emission filter prior to collection by a sCMOS camera. *Brightfield imaging*: The Orion system utilizes LED transillumination of the sample on the microscope slide. Chromogenic stains in the sample absorb a portion of the light, and the remainder is collected by the objective lens. The light follows the same path as the fluorescence emission described above, with the exception that a window is used instead of an emission filter. **e,** Validation of minimal channel crosstalk in 18-plex tonsil image after spectral extraction. Pearson's correlation coefficients between all channel pairs were calculated using the paired pixel intensities. Square boxes with colored borders denote excitation lasers. High correlation coefficients were only found in channel pairs that contains target markers that are in close proximity. **f.** Images of H&E-stained sections of colorectal cancer performed before IF imaging (0 cycles) and after one cycle of IF imaging (1 IF cycle) showing excellent preservation of staining intensity and morphology. Scalebars 5 mm.

#### Extended Data Figure 2

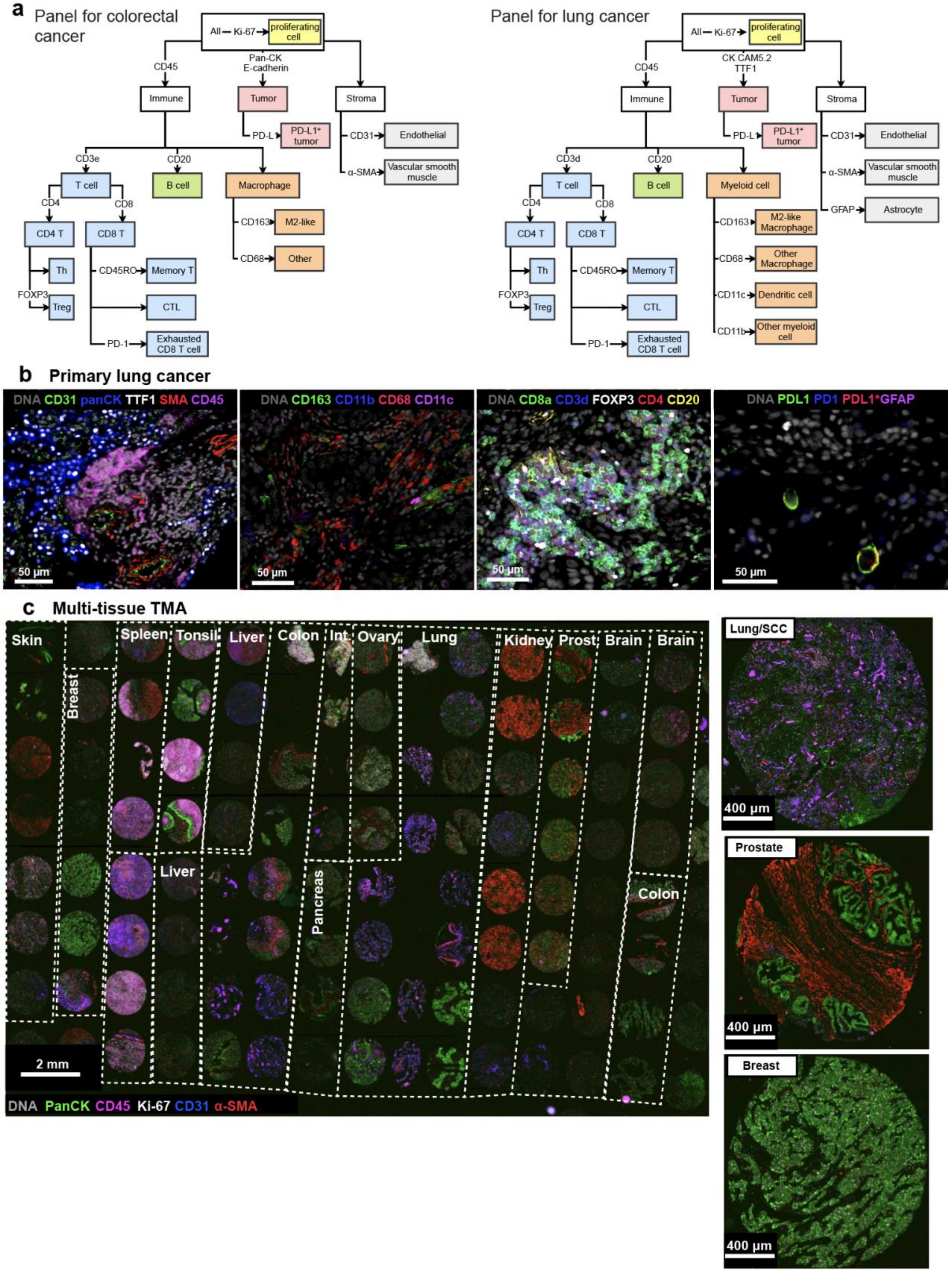


1166

**Extended Data Fig. 2 | Cell type calling for colorectal cancer and lung cancer and Orion imaging of different cancer histologies.**

**a,** Cell type calling dendrograms for Orion image analysis for colorectal cancer (left) and for lung cancer and lung cancer (right). First, immune, tumor, and stroma cells were identified using the indicated markers. Then, each of these classes of cells was further subclassified using the markers as noted. Ki-67 was used to distinguish proliferative from non-proliferative cells for each cell type. **b,** Representative images of 20-plex Orion panel from a primary lung adenocarcinoma sample. Note: two PD-L1 antibodies were used, PD-L1 (green) is E1L3N clone from Cell Signaling and PD-L1*(red) is EPR19759 from Abcam. Scalebars 50 μm. **c,** 16-plex (18 channel) Orion image from a tissue microarray (TMA) containing normal and diseased human tissues including inflammatory and neoplastic diseases (Examples highlighted are lung squamous cell carcinoma (SCC), prostate adenocarcinoma, and breast ductal carcinoma); DNA, pan-cytokeratin, Ki-67, α-SMA, CD45 and CD31 are displayed. Scalebars 2 mm and 400 μm, as indicated.

Extended Data Figure 3

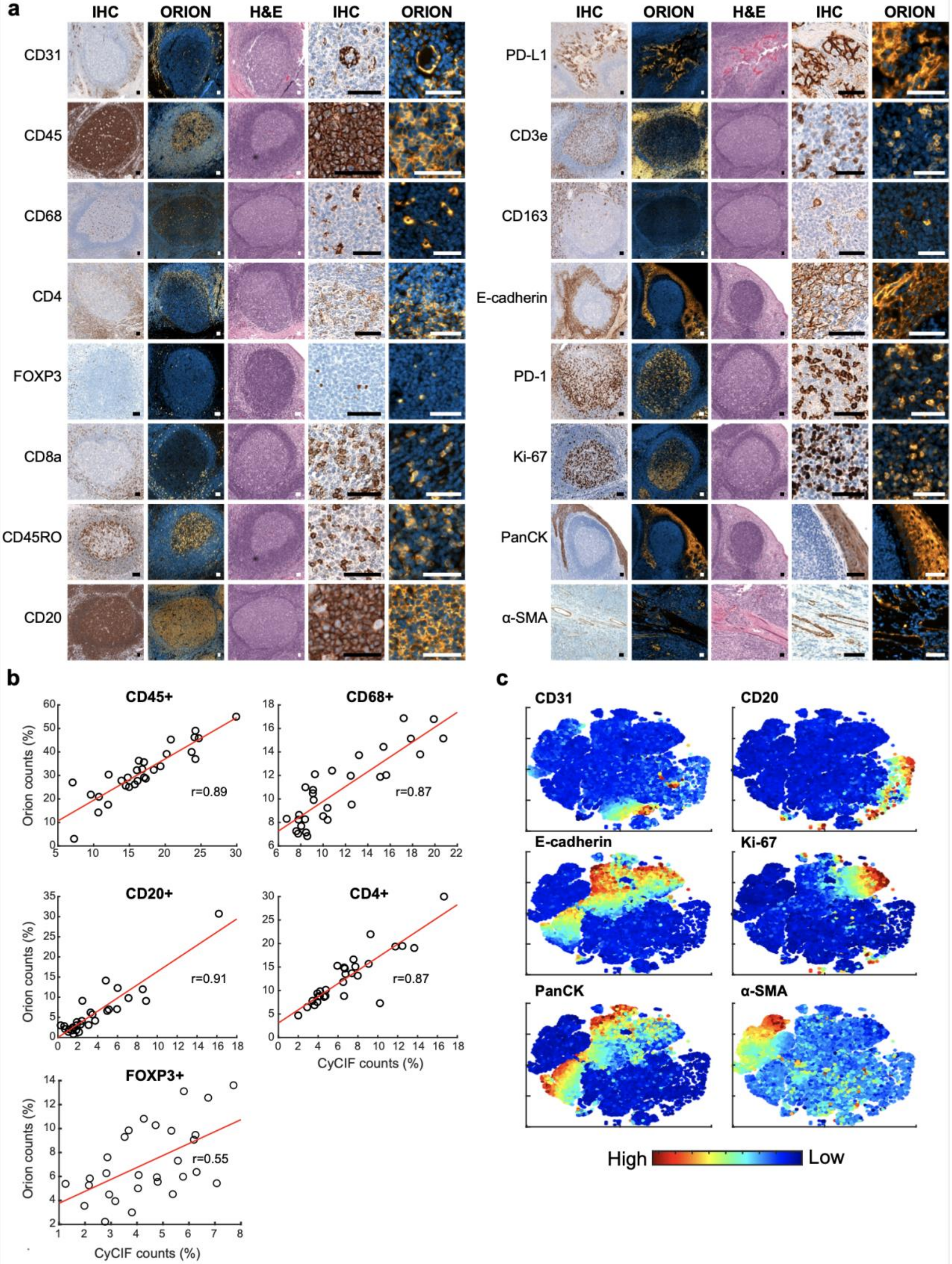


1181

**Extended Data Fig. 3 | Qualifying 16-plex single-shot Orion antibody panel relative to immunohistochemistry and Cyclic Immunofluorescence (CyCIF).**

**a,** Panels of images from FFPE tonsil sections showing single-antibody immunohistochemistry (IHC) for the indicated markers and matching channels extracted from the 16-plex Orion immunofluorescence (IF) images (H&E stain was performed on the same section as the Orion imaging). Scalebars 50 µm. **b,** Plots of the fraction of positive for the indicated markers (CD45, CD68, CD20, CD4, FOXP3) from whole slide Orion IF and CyCIF images acquired from two adjacent sections of 29 FFPE colorectal cancer specimens. Pearson correlation coefficients are indicated. **c,** t-distributed stochastic neighbor embedding (t-SNE) plots of cells from Orion IF image (specimen: C01). Log transformed marker intensities (CD31, CD20, E-cadherin, Ki-67, pan-cytokeratin, α-SMA) were used to color the dots in each panel.

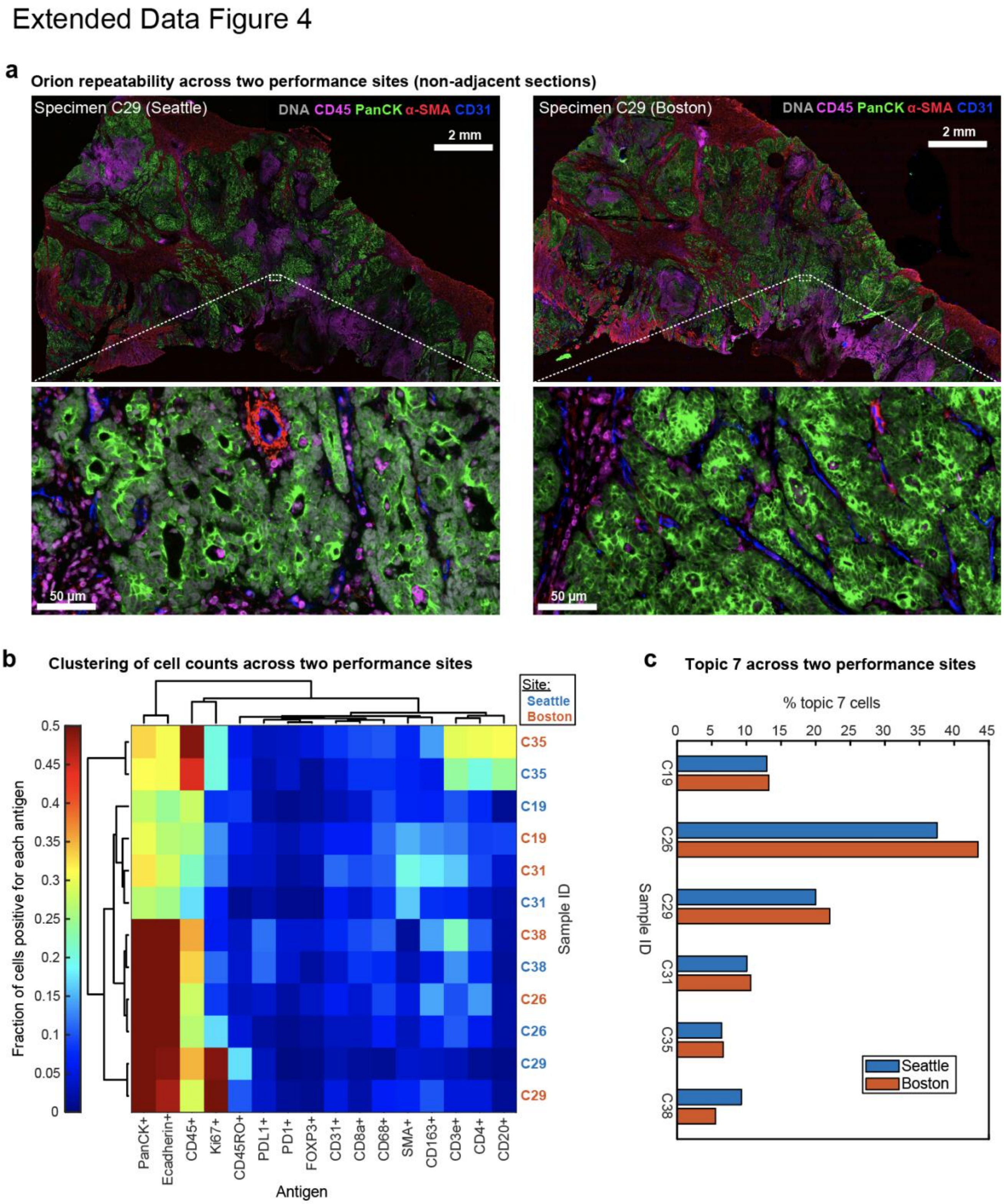


**Extended Data Fig. 4 | Evaluation of Orion data collected at two performance sites.**

**a,** Orion images of adjacent sections of sample C29 acquired in two different laboratories. Specimens on the left were imaged at RareCyte, Inc in Seattle WA and those on the right at HMS in Boston MA. DNA (Sytox), CD45, pan-cytokeratin, α-SMA, and CD31 are shown. Scalebars 2 mm and 50 µm, as indicated. **b,** Two-way hierarchical clustering heat map for the indicated markers and samples imaged at RareCyte (C19, C26, C29, C31, C35, C38) or at HMS (C19new, C26new, C29new, C31new, C35new, C38new) with the fraction of positive cells mapped to color. **c,** Bar plots showing the percentage of Topic 7 present in the indicated samples (C19, C26, C29, C31, C35, C38) imaged at two performance sites.

Extended Data Figure 5

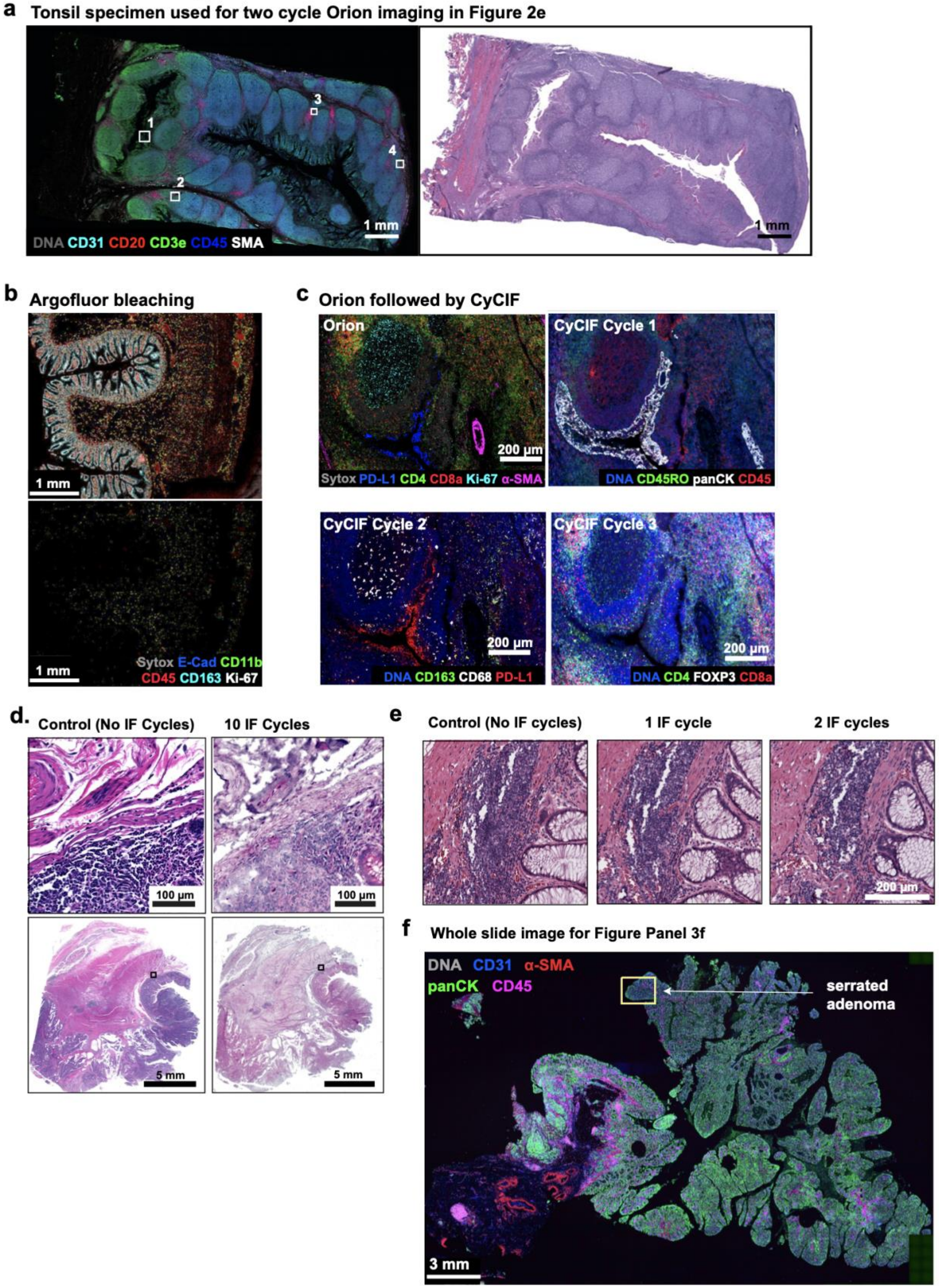


1204

**Extended Data Fig. 5 | Immunofluorescence and H&E images following multiple cycles of Orion imaging.**

**a,** Left panel: Orion image of FFPE tonsil showing DNA (Sytox), CD31, CD20, CD3e, CD45 and α-SMA). Scalebars 1 mm. ROIs 1 to 4 displayed in **Fig. 2e** are noted. Scalebar 50 µm. Right panel: H&E image after two cycle Orion imaging (i.e., after imaging of one panel, inactivation, and imaging of a second panel). Scalebars 1 mm. **b,** Top panel: Orion image of normal colon showing E-cadherin, CD11b, CD45, CD163, Ki-67, and DNA (Sytox) signal. Lower panel: same area of normal colon following inactivation of Orion fluorophores (see Methods). **c,** Same-slide Orion and CyCIF experiment. The tonsil samples were first processed with 16 Orion antibodies; PD-L1, CD4, CD8a, Ki-67, and α-SMA are shown. After imaging, fluorophores were inactivated by bleaching using the standard CyCIF protocol, then three-cycles of four-channel CyCIF staining and imaging were performed using the indicated antibodies. **d,** Images of H&E-stained sections of colorectal cancer without prior IF staining (right) and following 10 cycles of IF (left) using the standard CyCIF approach. Area shown in insets is indicated in the low magnification images. Scalebars 5 mm and 100 µm. **e,** Images of H&E-stained sections of colorectal cancer performed before IF imaging (0 cycles), after one cycle of IF imaging (1 IF cycle), and after two cycles of IF imaging (2 IF cycles). Scalebars 200 µm. **f,** Orion IF image from colorectal cancer resection specimen C26, showing an area of serrated adenoma with low pan-cytokeratin expression (markers as indicated). Higher magnification inset as indicated by the box is shown in **Fig. 3f.** Scalebar 3 mm.

Extended Data Figure 6

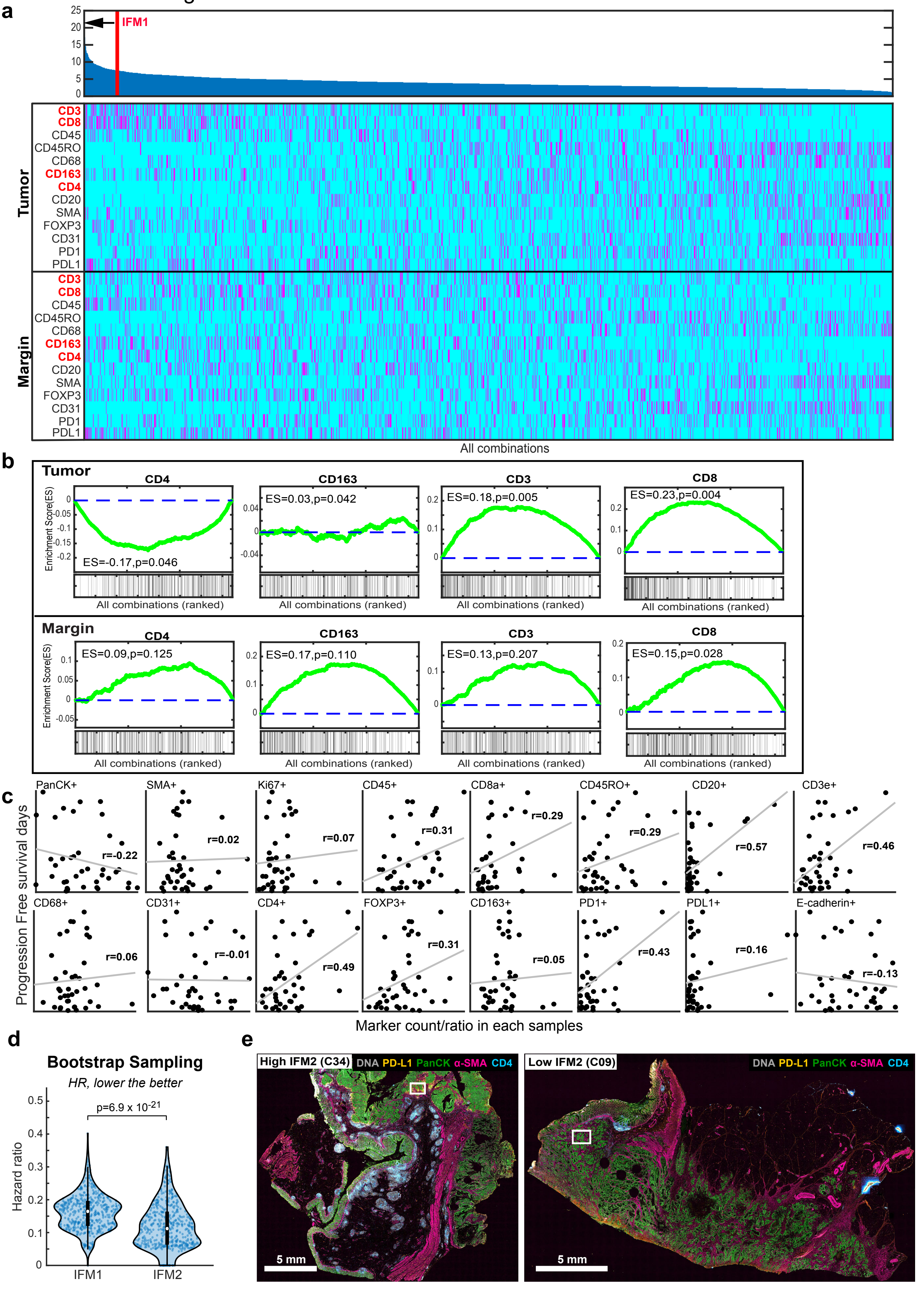

**Extended Data Fig. 6 | Assessment of individual markers in Image Feature Models of patient prognosis derived from Orion immunofluorescence images.**

**a,** Upper: Ranking of 1/hazard ratio (HR) for each Image Feature Model (IFM1 to IFM14,950) calculated by determining the frequency of cells positive for one or more of 13 markers in Orion IF images lying within (tumor center: CT) or outside of a region 100 μm from the tumor invasive margin (IM) model (n = 40 patients). Ranking position of IFM1 is indicated. IFM2 showed an HR = 0.08 (95% CI: 0.04-0.17, p = 1.91 x$10^{-06}$). Lower: Full heat map showing the selected markers at the tumor or margin in each combination. A total 14,950 combinations were generated as the set of 4 out of 26 parameters (13 markers in 2 regions). **b,** Enrichment plots showing enrichment scores (ES) for positive cells denoted by the indicated markers (and their location in the tumor or at the tumor margin) based on the 16-plex Orion images, indicating whether the marker/location feature is enriched in the Image Feature Models linked to the best hazard ratios. The green lines represent the running ES for a given marker/location as the analysis proceeds down the ranked list. The value at the peak is the final ES. **c,** Regression line scatter plot showing fraction of positive cells for indicated markers from the Orion 16-plex images vs. progression-free survival (PFS, days) for 40 patients with colorectal cancer. Each dot represents measurements from a single patient. $R^2$ for each plot is displayed. **d,** Plot bootstrapping of hazard ratios from IFM1 and IFM2 (unadjusted p = 4.62 x $10^{-26}$ and adjusted p = 6.9 x $10^{-21}$). Related to **Fig. 4f**. Detailed analysis is described in the methods section and pairwise two-tailed t-test were used unless otherwise mentioned. **e,** Representative Orion IF images of cases with high IFM2 (IS = 4 in specimen C34) and low IFM2 (IS = 0 in specimen C09). IF images show DNA, PD-L1, pan-cytokeratin, α-SMA, and CD4; Scalebars 5 mm. Higher magnification regions of interest are shown in **Fig. 4h**.

### Extended Data Figure 7

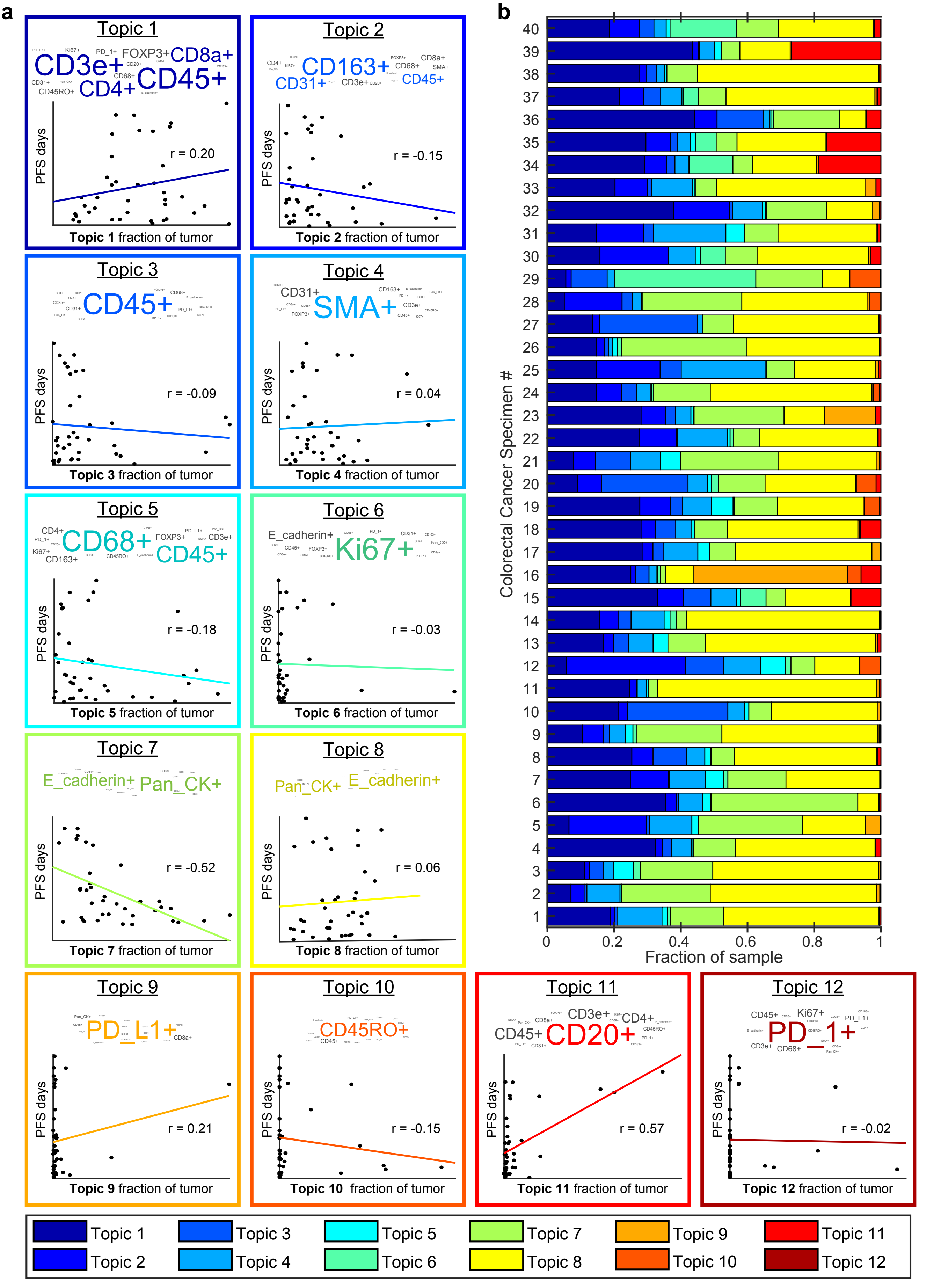

**Extended Data Fig. 7 | Cellular neighborhoods in colorectal cancer resections.**

**a,** Latent Dirichlet Allocation (LDA) probabilistic modeling was used to analyze Orion immunofluorescence data from 40 colorectal cancer specimens to reduce cell populations into neighborhoods ("topics") defined by patterns of single-cell marker expression. The analysis identified 12 topics that recurred across the dataset. Within each box is the LDA plot for the indicated topic (top) and a regression line scatter plot indicating the fraction of each tumor composed of the indicated LDA topic and the relationship to progression-free survival (PFS, days). Each dot represents measurements from a single patient. r value for each plot is displayed. **b,** Bar plot depicting the proportional distribution of the LDA Topics in the 40 colorectal cancer specimens.

Extended Data Figure 8

**a**

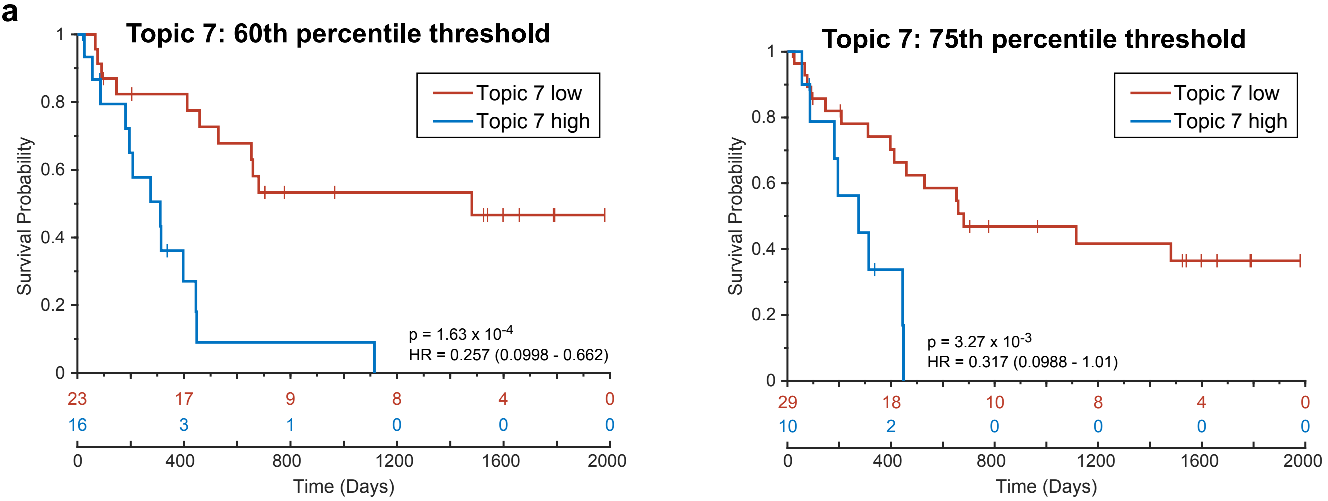


**b**

Ground truth (columns) vs. Prediction (rows)

| Prediction | Topic 0 | Topic 1 | Topic 2 | Topic 4 | Topic 7 | Topic 8 | Precision |
|---|---|---|---|---|---|---|---|
| Topic 0 | **0** 0.0% | **33** 0.3% | **8** 0.1% | **6** 0.1% | **1** 0.0% | **20** 0.2% | 0.0% 100% |
| Topic 1 | **6** 0.1% | **569** 4.8% | **22** 0.2% | **33** 0.3% | **18** 0.2% | **252** 2.1% | 63.2% 36.8% |
| Topic 2 | **635** 5.4% | **815** 6.9% | **1051** 8.9% | **1163** 9.9% | **378** 3.2% | **294** 2.5% | 24.2% 75.8% |
| Topic 4 | **15** 0.1% | **46** 0.4% | **70** 0.6% | **313** 2.7% | **24** 0.2% | **119** 1.0% | 53.3% 46.7% |
| Topic 7 | **214** 1.8% | **271** 2.3% | **69** 0.6% | **226** 1.9% | **2082** 17.7% | **960** 8.1% | 54.5% 45.5% |
| Topic 8 | **0** 0.0% | **180** 1.5% | **1** 0.0% | **0** 0.0% | **3** 0.0% | **1887** 16.0% | 91.1% 8.9% |
| Recall | 0.0% 100% | 29.7% 70.3% | 86.1% 13.9% | 18.0% 82.0% | 83.1% 16.9% | 53.4% 46.6% | **50.1% 49.9%** |

**c**

**Topic 8**

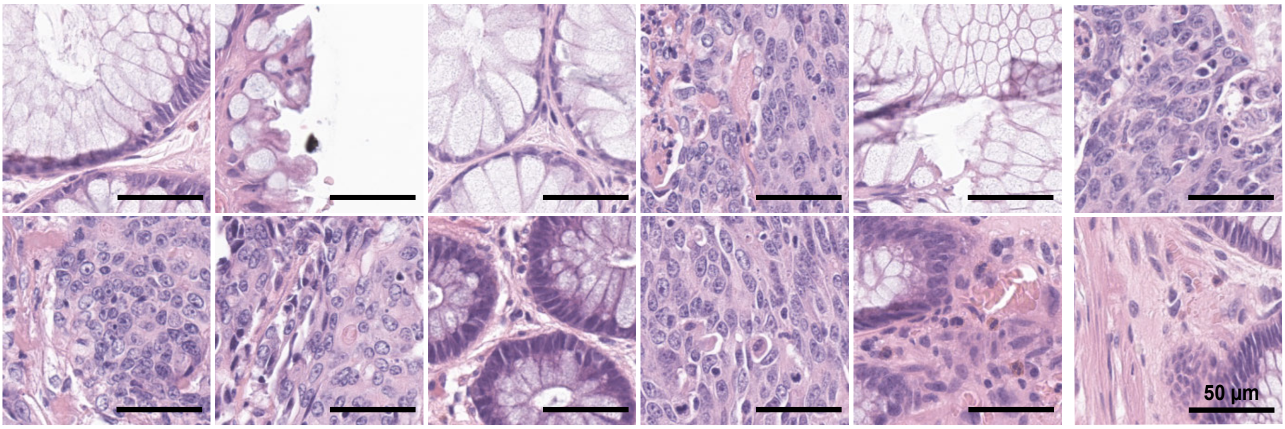

**Extended Data Fig. 8 | Evaluation of the performance of a Convolution Neural Network used to identify cellular neighborhood Topic 7 from H&E images of colorectal cancer.**

**a,** Kaplan Meier plots of PFS for 40 CRC patients based on the fraction of Topic 7 present in the tumor domain and stratified using a threshold ('cutoff') of $60^{th}$ percentile (left) and $75^{th}$ percentile (right) (HR, hazards ratio; 95% confidence interval; logrank p-value). **b,** Confusion matrix table showing performance of GoogLeNet convolutional neural network (CNN) trained using H&E data from Latent Dirichlet Allocation (LDA) Topic 7 and its performance in identifying Topic 7 cells from H&E data. Topic 0 contains the rest of the topics (3, 5, 6, 9, 10, 11, 12). Target class (ground truth) was assigned from LDA analysis of Orion images and Output class (predicted) was assigned by the GoogLeNet CNN. **c**, Gallery of representative H&E images of true positives for topic 8; Scalebars 50 µm.

Extended Data Figure 9

**CyCIF images of a CRC tumor rich in Topic 7 cells**

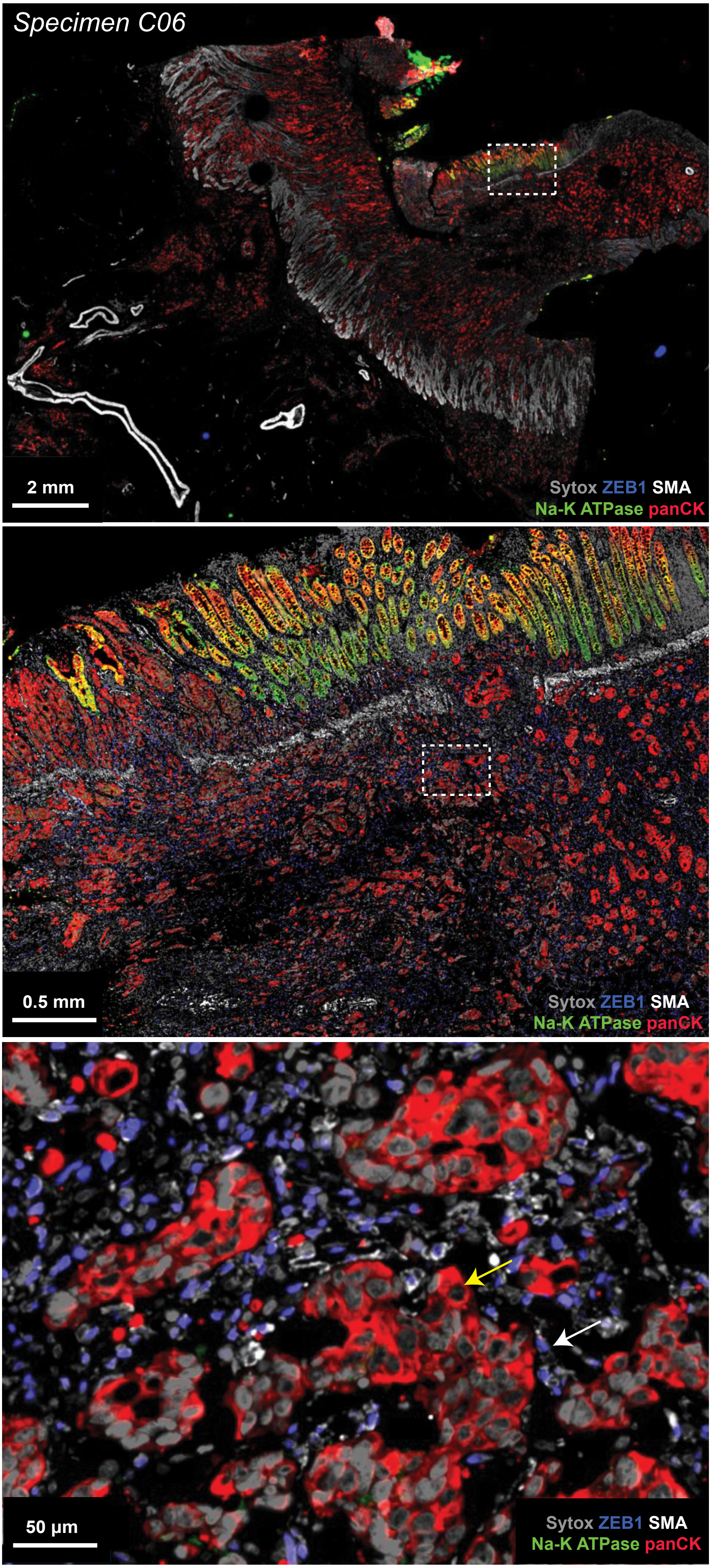

**Extended Data Fig. 9 | CyCIF imaging of Topic 7 tumor cells.**

CyCIF imaging of regions of specimen C06 which had a high fraction of Topic 7 cells. The CyCIF image is from a non-adjacent section to that used for Orion data section. Images show DNA (Sytox), ZEB1, α-SMA, NA-K ATPase, pan-cytokeratin. Location of insets are indicated. Scalebars, 2 mm, 0.5 mm, 50 µm, as indicated. The mesenchymal differentiation/EMT-marker ZEB1 (nuclear blue signal) is present in stromal cells (white arrow) but absent in the tumor cells (marked by pan-cytokeratin, red; yellow arrow).
